# Long noncoding RNA *HSCHARME* is altered in human cardiomyopathies and promotes stem cell-derived cardiomyocyte differentiation by splicing regulation

**DOI:** 10.1101/2024.11.06.622297

**Authors:** Giulia Buonaiuto, Fabio Desideri, Adriano Setti, Alessandro Palma, Angelo D’Angelo, Giulio Storari, Tiziana Santini, Pietro Laneve, Daniela Trisciuoglio, Monica Ballarino

**Affiliations:** Department of Biology and Biotechnologies “Charles Darwin”, Sapienza University of Rome, Rome, Italy; Center for Life Nano- & Neuro-Science of Istituto Italiano di Tecnologia (IIT), 00161, Rome, Italy; Institute of Molecular Biology and Pathology IBPM-CNR, 00185, Rome, Italy; Center for Molecular Biology Severo Ochoa (CBM Severo Ochoa) CSIC/UAM, Madrid 28049, Spain

**Keywords:** NcRNA, LncRNA, Splicing, hiPSCs, Cardiomyocytes, Cardiomyopathies

## Abstract

A growing body of evidence suggests that tissue-specific long noncoding RNAs (lncRNA) play pivotal roles in the heart. Here, we exploited the synteny between the mouse and human genomes to identify the novel lncRNA *HSCHARME* (*<u>H</u>uman <u>S</u>yntenic CHARME*) and combined single-cell transcriptomics, CAGE-seq data, RNA-FISH imaging and CRISPR-Cas9 genome editing to document its role in cardiomyogenesis. By investigating the mechanism of action of *HSCHARME* in hiPSC-derived cardiomyocytes, we found that the locus produces the major *pCHARME* isoform that associates with SC35-containing speckles and interacts with the splicing regulator PTBP1. Consistently, the functional inactivation of *pCHARME* influences the splicing of cardiac-specific pre-mRNAs and impacts their expression, which parallels a decline in cardiomyocyte differentiation and physiology. In line with a possible association with disease, large-scale analysis of the lncRNA expression across cardiomyopathy patients revealed increased levels of *pCHARME* in hypertrophic (HCM) and dilated (DCM) hearts and identified a subset of disease-associated targets whose expression can be modulated through *HSCHARME* dosage.

By unlocking mechanistic insights into the role of *pCHARME* in cardiac cells, our data identify a novel non-coding regulator of cardiomyocyte function with potential implications in disease.

## INTRODUCTION

Heart diseases represent the primary cause of death globally^1^. As the cardiac muscle fails to renew cardiomyocytes (CM)^2^ regenerative medicine therapies are considered crucial for stimulating the repair of damaged hearts, thereby reducing the need for transplants. Nonetheless, the restoration of fully matured and functional cardiomyocytes is still challenging due to the partial knowledge of the factors regulating their endogenous development and maturation^3^. Over the years, considerable attention has been placed on coding genes^4^. More recently, the increasing discovery of long-noncoding RNAs (lncRNAs) has revealed unexplored modalities that are essential for CM to acquire their identity and functionality^5,6^. LncRNAs represent a heterogeneous class of transcripts longer than 500 nucleotides, with limited protein-coding capability^7^. Since their discovery, they have been implicated in regulating every step of the gene life cycle^8^, as well as a wide range of processes, from physiological to pathological^9^. The remarkable temporal and tissue-specific expression of these molecules designates them as optimal regulators of physiological organ development. In the heart, several studies conducted in murine models have underscored the critical roles of lncRNAs in sustaining cardiac homeostasis, with their dysregulation frequently associated to the onset of disease^6,10–14^. Despite the presence of many orthologues, their direct investigations in human cells or tissues remain scarce. Nonetheless, several lncRNAs have been implicated in human CM processes, including proliferation and regeneration^5,15,16^.

In the nucleus, lncRNAs fine-tune gene expression by interacting with DNA/RNA molecules, RNA binding proteins, or by shaping the genome three-dimensional organization^17^. Their role as RNA scaffolds is exemplified by their ability to form RNA-rich condensates and create high-local molecule concentrations at specific loci. This is the case of *NEAT1* and *MALAT1*, which accumulate within the nucleus as RNA components of specific sub-nuclear compartments enriched in splicing factors, thereby regulating pre-mRNA splicing^18,19^. In mice, we have previously identified *pCharme*, an architectural lncRNA coordinating the activation of pro-myogenic genes at specific nuclear condensates^20,21^. In the heart, the genetic ablation of *pCharme* causes persistent expression of fetal-like gene programs delaying CM maturation, which ultimately leads to dilated cardiomyopathy^22^. This highlights an important role for the lncRNA in the pathophysiology of the heart and encourages further analyses of its possible contribution in humans. Built on this rationale, we used comparative genomics to further investigate conserved synteny at the previously identified *pCharme* locus^20^. We found that the human syntenic gene, *HSCHARME*, is expressed in the human heart, both in fetal and adult CM. By using cardiomyocytes derived from human induced pluripotent stem cells (hiPSC-CM), we show that the *pCHARME* transcript localises close to SC35-containing speckles and directly binds to the Polypyrimidine tract-binding protein 1 (PTBP1) to influence the splicing of cardiac-specific pre-mRNAs involved in CM differentiation. Finally, we found that the endogenous *pCHARME* expression is significantly increased in dilated (DCM) and hypertrophic (HCM) human cardiomyopathies, correlating with dysregulation of disease-associated genes. Overall, these results identify *pCHARME* as a new gene linked to cardiac disease, with potential implications for therapeutic approaches in human CM.

## RESULTS

### Identification of conserved synteny of the *pCharme* locus in *Homo sapiens*

To explore the phylogenetic conservation of the murine *pCharme*^20,21^ and precisely map the chromosomal location of the gene in the human genome assembly, we performed a cross-species analysis through the UCSC genome browser “LiftOver” tool (**Fig. 1A**). Conversion from the murine (mm10; chr7:44,473,538-44,486,138) into the human coordinates revealed the existence of a syntenic locus (hg38; chr19:50,486,482-50,500,386), herein named *HSCHARME* (gene aliases lncFAM^23^ and MYREM^24^), whose mapping located a 94-nucleotide extension at the beginning of the gene, as compared to the previous annotation^20^.

**Fig. 1.**
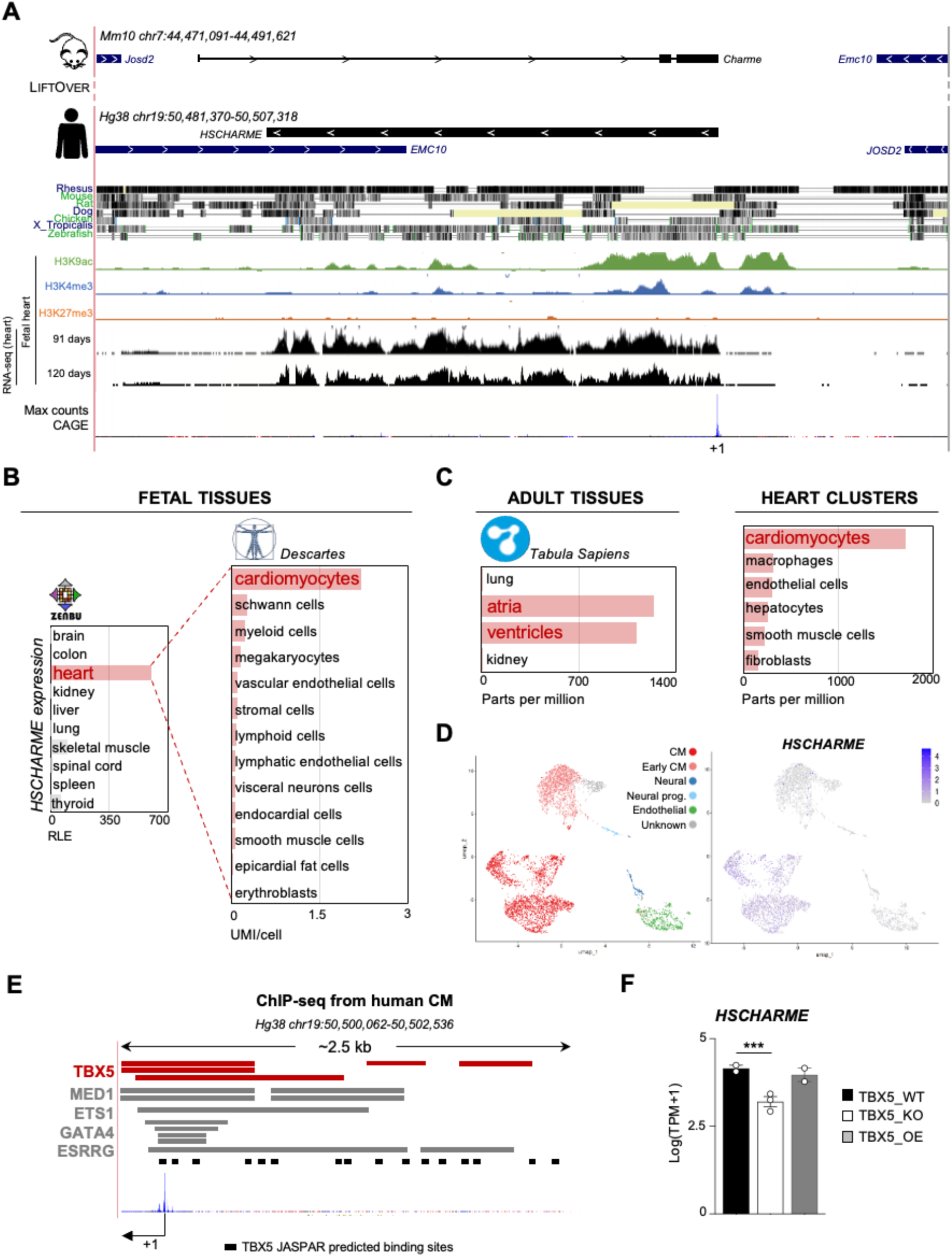
Identification of conserved synteny of the *pCharme* locus in Homo sapiens. **A)** UCSC visualization showing the chromosome position and the genomic coordinates of *Charme* gene in mice and humans (*HSCHARME*). UCSC default tracks for Vertebrate Multiz Alignment & Conservation and Histone modifications (H3K9ac, green; H3K4me3, blue; H3K27me3; orange) are also shown, together with the RNA-seq reads taken from GEO and FANTOM5 CAGE datasets. TSS= transcriptional start site. **B)** Left panel: quantification of *HSCHARME* TSS usage (ROI coordinates: Hg38 chr19:50,476,155-50,510,700) in fetal tissues from CAGE data (Phase1 and 2 datasets) from ZENBU FANTOM5 Human hg19 promoterome. For each specific sample, bars represent the Relative Logaritmic Expression (RLE) of the Tag Per Million values of TSS usage. Right panel: quantification *HSCHARME* expression in cardiac cells subtypes from Descartes scRNA-seq atlas27. Bars represent UMI/cell values. **C)** Quantification of *HSCHARME* expression in adult tissues from scRNA-seq data of Tabula Sapiens28 organized by full details (left panel) or by tissue and cell type (right panel) showing *HSCHARME* expression. Bars represent the Parts Per Million values of *HSCHARME* TSS usage per sample. **D)** Left Panel: Integrated UMAP plot of single-cell transcriptomic profiles from hiPSC-CM (2857 cells from Day 14, 2321 cells from Day 45, as in ^31^,describing cell identity assignment to CM, Early CM, Neural, Neural progenitors, Endothelial, or Unknown (cells that could not be assigned to any specific identity) subpopulations. Right panel: *HSCHARME* expression at single-cell resolution over the UMAP representation of D14 and D45 hiPSC-CM dataset. **E)** UCSC visualization of ReMap ChIP-seq track across *HSCHARME* promoter in human CM. Analysis was performed by restricting the biotype selection to CM and focusing on the 2.5 Kb region upstream *HSCHARME* locus. The genomic coordinates, the *HSCHARME* TSS (+1, black arrow) and the TBX5 JASPAR binding sites (black squares) predicted with JASPAR 2022 (29) (relative profile score threshold=80%) are shown. **F)** Quantification of *HSCHARME* expression from RNA-seq analysis performed in n=2 wild type (WT), n=3 TBX5 knockout (KO) and n=2 TBX5 overexpressing (OE) hiPSC-derived CM (GSE8158530). Data are represented as Log(TPM+1) to avoid negative values with TPM= Transcript per Million mapped reads. Data information: ***p < 0.001, adjusted p-value.

As shown in **Fig. 1A**, we found an overall high-level sequence conservation of the ∼26 kb-long region, especially in Rhesus, Mouse, X. Tropicalis and Zebrafish, as well as the synteny of the *EMC10* and *JOSD2* nearby coding genes. More specifically, sequence conservation analysis between the mouse and human genes (intronic and exonic regions) highlighted a degree of 45.1 % of sequence identity (**Supplementary Fig. 1A**), which positions *HSCHARME* between the range of well-known lncRNAs, such as *NEAT1* and *XIST*^25,26^ (**Supplementary Fig. 1B**). Consistent with the transcriptional activity of the locus, Transcriptional Start Site (TSS) mapping by CAGE (Cap Analysis of Gene Expression) confirmed the presence of a sharp peak on the negative strand of the *HSCHARME* gene, attributable to the existence of a transcript produced in antisense direction (**Fig. 1A**).

We then searched available datasets for epigenetic and gene expression signatures across the human gene focusing on cardiac samples. We found that embryonal whole-heart samples show deposition of histone H3 acetyl-lysine 9 (H3K9ac; GSM706849^27^), trimethylation of histone H3 lysine 4 (H3K4me3; GSM772735^27^) and the absence of the repressive trimethylation of histone H3 lysine 27 (H3K27me3; GSM621450^27^) marks, correlating with the transcriptional activation of the locus in fetal hearts (90-120 days), as further confirmed by the gene expression RNA-seq outputs (GSM1059494; GSM1059495) (**Fig. 1A**). CAGE-sequencing data from the FANTOM5 human promoterome catalogue^28^ (https://fantom.gsc.riken.jp/zenbu), further confirmed the specific expression of *HSCHARME* in both fetal hearts (84-217 days) and skeletal muscle cells (**Fig. 1B**, left panel). In line with our previous finding in mice^22^, scRNA-seq data from Descartes^29^ and Tabula Sapiens^30^ atlases revealed the highest expression of *HSCHARME* in fetal (72–129 days) cardiomyocytes (**Fig. 1B**, right panel). Restriction of lncRNA expression to cardiomyocytes persists in adult hearts, with no evident difference between atria and ventricles (**Fig. 1C**, left and right panels). Further examination of scRNA-seq data derived from hiPSC differentiated into cardiac cells^31^ definitively confirmed the highly specific expression of *HSCHARME* within the CM cluster (**Fig. 1D**, **Supplementary Fig. 1C-E** and **Supplementary Table 1,** see **Materials and Methods** for details).

Searching for possible regulators of *HSCHARME* expression we mined ReMap ChIP-seq atlas and found the binding of well-known transcription factors in the 2.5 Kb region from the TSS (**Fig. 1E**) with only *TBX5* (T-box transcription factor) showing clear and specific expression in CM (**Supplementary Fig. 1F**). Consistent with the functional implication of TBX5 in *HSCHARME* expression, computational analyses of available transcriptomic datasets from WT, TBX5-KO and TBX5-OE hiPSC-derived CM^31^, demonstrate a significant decrease of the lncRNA upon the loss-of-function of TBX5 (**Fig. 1F**). Along with our previous findings in mice^22^, these results underscore a conserved role for TBX5 in positively regulating the expression of *HSCHARME* in the human heart. Since the overexpression of TBX5 alone does not influence the levels of the lncRNA, we argue that it may establish its basal transcriptional expression, while other factors could be involved in the transcriptional regulation of the locus in cardiac cells.

Overall, these new data provide a high-resolution map of *HSCHARME* expression in the human heart and detail the cell-type specific restriction of the lncRNA to CM. The similarities of *HSCHARME* with its murine counterpart, which has already been shown to play an important role in cardiac remodeling^20–22^, encouraged further investigations into the relevance of the human transcript in disease.

### *HSCHARME* characterization in hiPSC-derived human cardiomyocytes

Considering the specific expression of *HSCHARME* in CM, we used hiPSCs as a model to study cardiomyogenic differentiation. The applied protocol^32^ exploits the induction of the WNT/β-catenin pathway (Day 0, CHIR addition), to guide cells towards mesoderm followed by its subsequent inhibition (Day 3, IWR-1 addition) to promote a CM fate (**Fig. 2A**). In line with the acquisition of the CM identity, RT-qPCR analyses performed at specific time points of hiPSC differentiation show the expected dynamic wave of cardiac gene expression^32^ (**Fig. 2B**), alongside the onset of spontaneous beating of cellular foci, starting at days 8-10 (**Supplementary Video 1**). The expression of *HSCHARME* at day 10 and its progressive increase (**Fig. 2B**) suggest its potential association with human CM commitment and differentiation. To directly test this hypothesis, we used a CRISPR/Cas9-based gene editing technology to generate *HSCHARME* knock-out (KO) hiPSCs. Specifically, sgRNAs were designed to produce a genomic deletion (∼4.8 Kb sized) overlapping *HSCHARME* TSS, which also includes the TBX5 binding sites (delta promoter=ΔP) (**Fig. 2C**). Mutant hiPSCs-CM were checked to confirm the correct genomic editing, the abrogation of *HSCHARME* expression (**Supplementary Fig. 2A**) and the absence of possible off-targets (**Supplementary Fig. 2B**, **Supplementary File 1** and **Supplementary Table 2)**. Following these verifications, we proceeded with the transcriptome analysis of WT *versus* isogenic ΔP-CM.

**Fig. 2.**
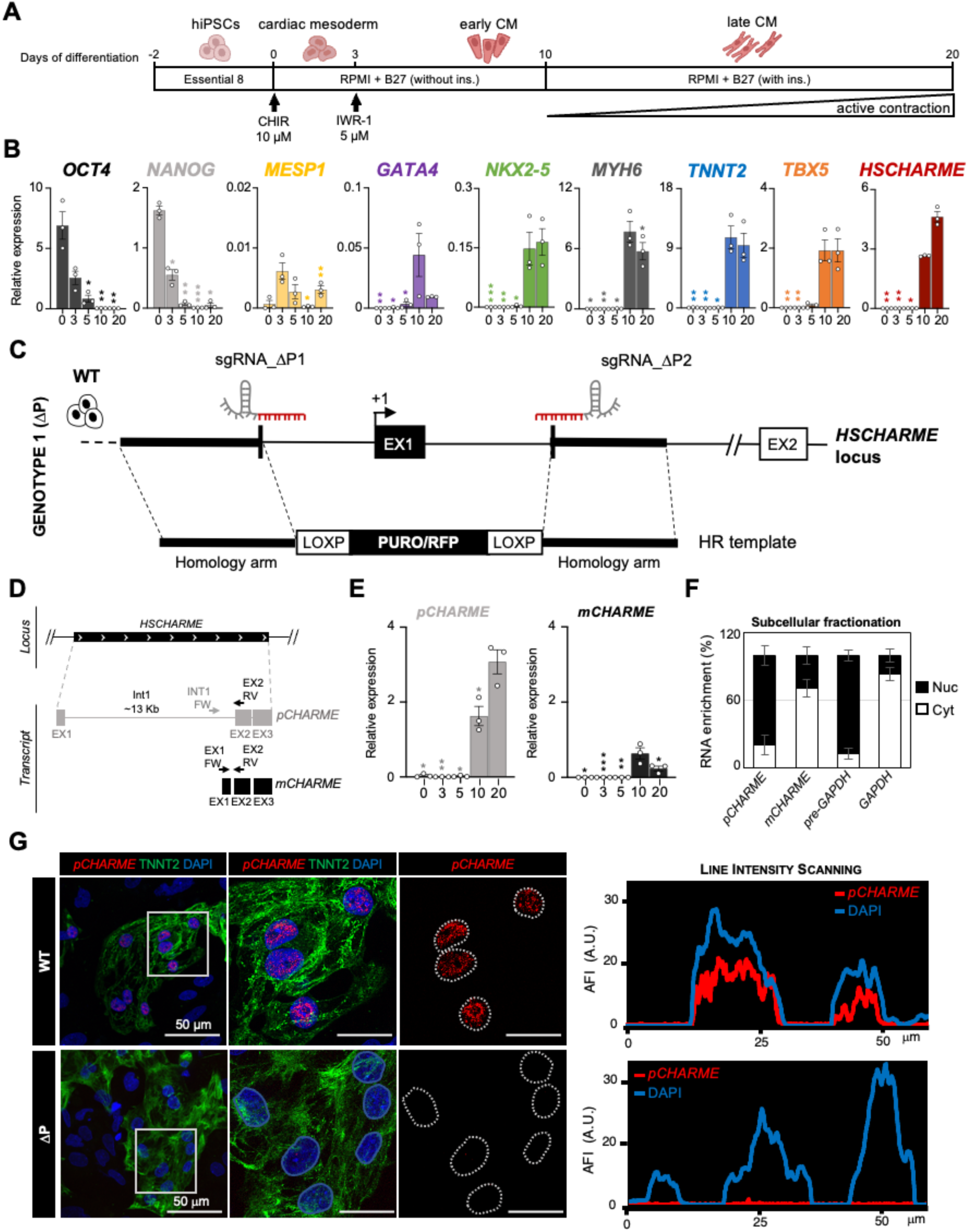
*HSCHARME* characterization in hiPSC-derived human CM. **A)** Schematic representation of CM differentiation from hiPSCs. **B)** RT-qPCR amplification of readout mRNAs as well as *HSCHARME* from total RNA extracted from hiPSCs-derived CM at specific time points (D0-D20). Data were normalized to *ATP5O* mRNA and represent relative expression means (2^^-DCt^) ± SEM of 3 biological replicates. Statistical tests were conducted on log₂-transformed fold change (LogFC) values compared to the control sample (the timepoint with the highest expression). **C)** Schematic representation of the genome editing strategy design followed to obtain ΔP hiPSC cell line using CRISPR/Cas9 technology. **D)** Schematic representation of *HSCHARME* isoforms as reconstructed by genome assembly. Arrows represent the position of primers for RT-qPCR. **E)** RT-qPCR amplification of *pCHARME* and *mCHARME* from total RNA extracted from hiPSC-derived CM at specific time points (D0-D20). Data were normalized to *ATP5O* mRNA and represent relative expression means (2^^-DCt^) ± SEM of 3 biological experiments. Statistical tests were conducted on log₂-transformed fold change (LogFC) values compared to the control sample (the timepoint with the highest expression). **F)** Quantification of the subcellular distribution of *pCHARME* and *mCHARME* isoforms from biochemical fractionation of D10 CM. The histogram shows the RT-qPCR quantification of the relative % of RNA abundance in cytoplasmic versus nuclear compartments. *GAPDH* and *pre-GAPDH* RNAs were used, respectively, as cytoplasmic and nuclear controls. **G)** Left panel: Representative RNA-FISH staining for *pCHARME* (red) combined with TNNT2 immunofluorescence (green) in WT and ΔP hiPSC-derived D20 CM. Panels represent the maximun 2D projection of full-size image of confocal caption, digital magnification of region highlighted by white squares and *pCHARME* signal distribution inside the nuclei highlighted by dotted line. Right panel: plot displaying Average Fluorescence Intensity (AFI) of *pCHARME* RNA-FISH and DAPI signals of a single focal plane. Overlapping of the lines indicates colocalization between the signals scanned along the horizontal distance of the selection (white rectangle). Data information: *p < 0.05, **p < 0.01, ***p < 0.001; one-sample two-tailed Student’s t-test against the null hypothesis of zero (no change).

*De novo* assembly of reads from the WT condition suggested that two *HSCHARME* full-length isoforms are produced in CM, the partially spliced transcript (precursor) *pCHARME* stably retaining the first intron and the fully spliced transcript (mature) *mCHARME* (**Fig. 2D** and **Supplementary Fig. 2C**). Both isoforms were induced throughout CM differentiation, with *pCHARME* being consistently more abundant than *mCHARME* and enriched to nuclei (**Fig. 2E-F**). Subcellular fractionation assays followed by RT-qPCR analyses revealed a cytoplasmic enrichment for *mCHARME.* The presence of open reading frames within *mCHARME*, possibly translated into micro-peptides^33–36^, was excluded by the Coding Potential Calculator tool (CPC2, http://cpc2.gao-lab.org/index.php). In line with the biochemical fractionation, high-resolution RNA-fluorescence *in situ* hybridization (RNA-FISH) experiments using probes against *pCHARME* confirmed its nuclear localisation (**Fig. 2G**). No fluorescence was detected in ΔP-CM, which proves the specificity of *pCHARME* signals and the efficiency of the lncRNA KO. The latter was further confirmed by i) RT-qPCR analysis of WT and ΔP-CM with *pCHARME* and *mCHARME* specific primers (**Fig. 3A**, middle panel), ii) TPM quantification of the transcriptomic reads and ii) the RNA-seq genomic plots (**Fig. 3A**, right panel and **Supplementary Fig. 2D**).

**Fig. 3.**
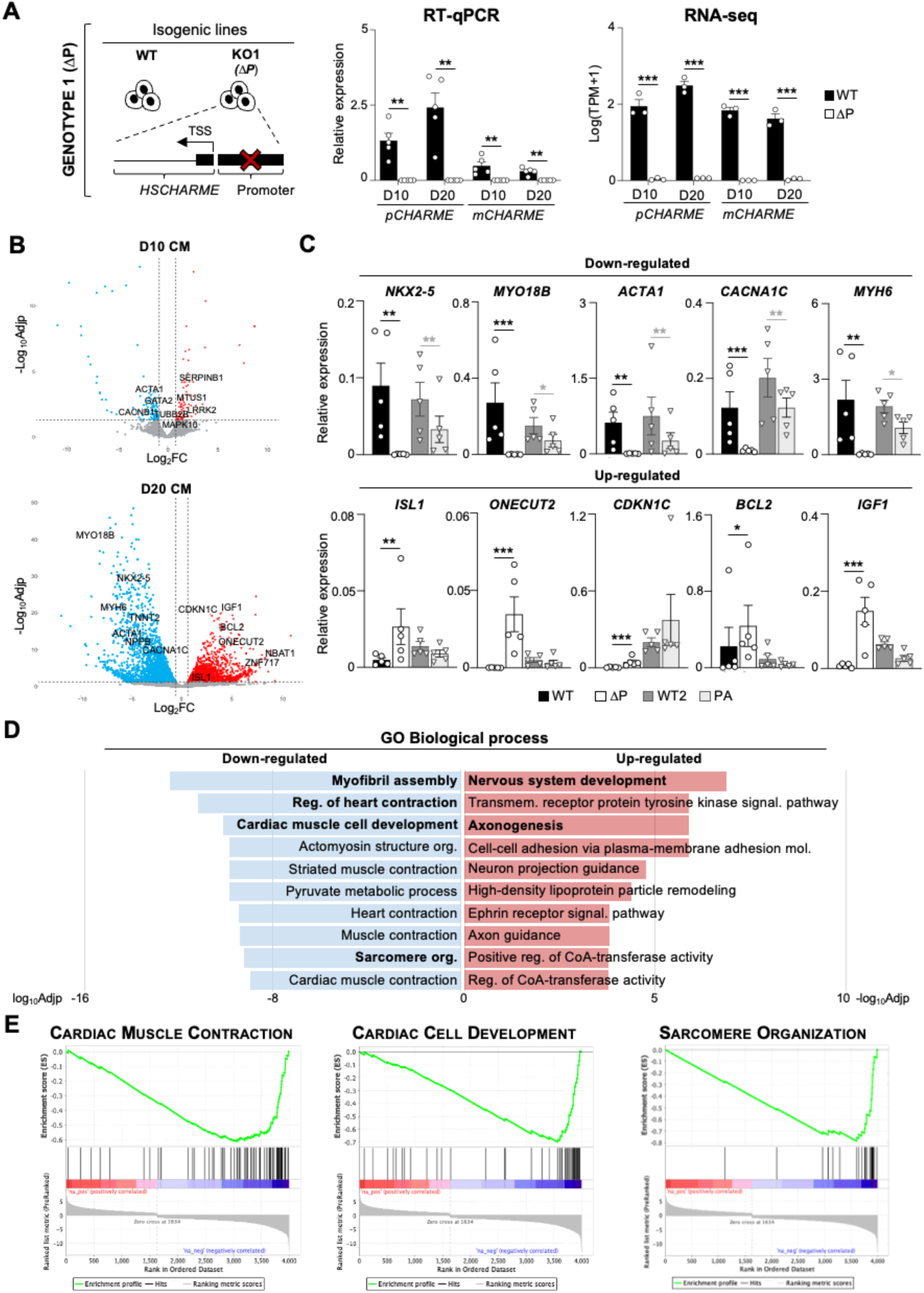
Genome-wide profiling of WT and ΔP hiPSC-derived CM. **A)** Left panel: Schematic representation of the genome editing strategy design followed to obtain ΔP hiPSC cell line using CRISPR/Cas9 technology. Middle panel: RT-qPCR amplification of *pCHARME* and *mCHARME* from WT vs ΔP D10 and D20 CM. Data were normalized to *ATP5O* mRNA and represent relative expression means (2^^-DCt^) ± SEM of 5 biological replicates. Data information: **p < 0.01; one-sample two-tailed Student’s t-test against the null hypothesis of zero (no change). Right panel: Quantification by RNA-seq (TPM) of *pCHARME* and *mCHARME* expression in WT vs ΔP hiPSC-derived D10 and D20 CM. Data are represented as Log(TPM+1) to avoid negative values. Data information: ***p < 0.001, FDR. **B)** Volcano plots showing differentially expressed genes (DEGs) from the RNA-seq analysis of WT vs ΔP hiPSC-derived CM at D10 vs CTRL at the same timepoint (upper panel) and D20 vs CTRL at the same timepoint (lower panel). Significantly up-regulated (FDR<0.05; log_2_FC>1) and down-regulated (FDR<0.05; log_2_FC<-1) genes are represented in red and blue respectively. **C)** RT-qPCR quantification of down-regulated (upper panel) and up-regulated (lower panel) DEGs in WT vs ΔP (black) and WT2 vs PA (grey) hiPSC-derived D20 CM. Data were normalized to *ATP5O* mRNA and represent relative expression means (2^^-DCt^) ± SEM of 5 biological experiments. Statistical tests were conducted on log₂-transformed fold change (LogFC) values compared to the control sample (WT for ΔP-CM and WT2 for PA-CM). Data information: *p < 0.05, **p < 0.01, ***p < 0.001; one-sample two-tailed Student’s t-test against the null hypothesis of zero (no change). **D)** Gene ontology (GO) enrichment analysis performed with EnrichR on down-regulated (left) and up-regulated (right) DEGs in WT vs ΔP hiPSC-derived D20 CM. Bars indicate +/–log10 adjusted p-value (log10Adjp and –log10Adjp) of the top enriched biological processes. All the represented categories show an Adjp<0.05. **E)** GSEA plot showing the enrichment of “cardiac muscle contraction”, “cardiac cell development” and “sarcomere organization” processes resulting down-regulated in WT vs ΔP D20 CM.

As previously observed for the murine orthologue^22^, the staining of *pCHARME* in human CM exhibits a discrete nuclear pattern. Nonetheless, here we noticed that the fluorescent signals encompass multiple foci, which suggests *trans*-regulatory roles for the lncRNA. In this direction, differential expression analysis performed to compare the WT and *Δ*P transcriptomic datasets, revealed an impact of *HSCHARME* on the expression of 175 and 3637 genes at 10 (D10) and 20 (D20) differentiation days, respectively (-1<log_2_FC>1; FDR<0.05) (**Fig. 3B**, **Supplementary Fig. 3A** and **Supplementary Table 3**). Of them, a total of 1465 differential expressed genes (DEGs) were up-regulated (n=68 at D10; n=1397 at D20), whereas 2347 were down-regulated (n=107 at D10; n=2240 at D20) in ΔP-CM. Importantly, none of the putative off-target genes was found among DEGs (**Supplementary Fig. 3B**), which conclusively excludes their involvement in the transcriptomic alterations. Moreover, we did not find alteration in the expression of *pCHARME* neighbouring loci (±260 Kb), which conclusively excludes a role for the lncRNA on nearby gene expression in cardiac cells (**Supplementary Fig. 3C**). To strength these results, we produced an independent knock-out cell line (PA-hiPSC), with a different *HSCHARME* mutation and genetic background (WTSIi004-A; referred to as WT2). Specifically, the PA-hiPSC were generated through the insertion of a strong termination cassette inside the *HSCHARME* locus using a strategy previously setup in mice^20^ (**Supplementary Fig. 3D**). Upon checking the correct abrogation of *pCHARME* and *mCHARME* (**Supplementary Fig. 3E**), the absence of off-target mutations was confirmed by DNA sequencing (**Supplementary Fig. 3F**, **Supplementary File 1** and **Supplementary Table 2**). To note, no off-target gene was found in common between the two genotypes, which excludes possible effects due to unwanted Cas9 background activities. Validation by RT-qPCR analyses performed on the top-most down-regulated DEGs from ΔP-CM showed their significant decrease in both the mutant cell lines (**Fig. 3C** and **Supplementary Fig. 3G**). As both CRISPR-Cas9 strategies demonstrate high efficiency in the *pCHARME* knockout (∼99%), the minor alterations in gene expression observed within the PA context can be attributed to the intrinsic features of the iPSC-WTSIi004-A background.

Gene Ontology (GO) term enrichment analysis performed on transcriptomic datasets revealed that at early stages of CM differentiation (D10), down-regulated DEGs are associated with developmental pathways, such as limb development (**Supplementary Fig. 3H**, left and **Supplementary Table 3**). At later stages (D20), enriched GO categories relate to CM structure and function, including myofibril assembly, heart contraction and cardiac muscle cell development gene classes (**Fig. 3D**, left and **Supplementary Table 3**). Gene set enrichment analysis (GSEA) confirmed the significant downregulation of genes belonging to these categories (**Fig. 3E**), with key examples that include *NKX2-5*, crucial for CM development^37^, along with structural and functional components such as *MYH6*^38^, *MYO18B*^39^, and *CACNA1C*^40^. Conversely, upregulated genes enriched GO categories relates to axonogenesis and nervous system development (**Fig. 3D**, right and **Supplementary Table 3**), may reflect the predisposition of the embryonic stem cells to differentiate toward default neuronal fate in the absence of additional stimuli^41^. We argued that *HSCHARME* depletion, by inhibiting the signals activating CM specification, might promote the upregulation of alternative pathways leading to a default neuronal state.

To assess the evolutionarily conservation of *pCHARME* function between human and mouse, significant DEGs from WT and *Charme* KO post-natal hearts (GSE200878^22^) were analyzed, identifying 847 human homologs (**Supplementary Fig. 3I**). Of them, 208 genes (24.55% of the murine DEGs) were commonly deregulated upon *pCHARME* ablation in cardiac muscle (i.e. *MYO18B*, *CACNA1C*, and *NPPB)*, enriched in pathways related to muscle homeostasis (e.g., glycolytic process, heart development), muscle function (e.g., contraction, fatty acid metabolism), and linked to cardiac disorders such as familial atrial fibrillation and hypertrophic cardiomyopathy. Collectively, the transcriptomic data, evolutionary conservation, and the functional interpretation of our datasets support the role of *HSCHARME* as a positive regulator of genes whose expression is physiologically relevant for the differentiation of CM.

### Functional implication of *HSCHARME* in cardiomyogenesis

To functionally characterize WT and gene-edited hiPSC-CM, we tested their capacity to contract. By tracking the contraction dynamics through MUSCLEMOTION (**Supplementary Fig. 4A**) we found a significant decrease in the beat rate of mutant ΔP-CM as well as the alteration of other parameters, such as time to peak, duration and peak to peak time (**Fig. 4A**). These changes are consistent with a general decrease of the spontaneous beating frequency in *Δ*P-CM compared to WT cells, with individual contractions becoming longer upon *pCHARME* loss. Importantly, the inspection of cells over differentiation evidenced a significant delay in the onset of beating of mutant CM (*Δ*P and PA) compared to their WT counterparts (11 *versus* 8 days on average, **Fig. 4B** and **Supplementary Fig. 4B**). Other phenotypical traits associated with cellular morphology^42^ were influenced by *pCHARME* loss-of-function with mutant CM appearing smaller and rounder than the isogenic WT (**Fig. 4C** and **Supplementary Fig. 4C**), indicating a possible delay in CM differentiation. To deepen our understanding of the timing of *pCHARME* regulation across differentiation, we performed flow cytometry analysis of PDGFRA^+^/CD56^+^ cardiac progenitors and PDGFRA^+^/CD82^+^ cardiac-committed progenitors which show that *pCHARME* loss leads to a 43% reduction in cardiac progenitors and 20% reduction in the cardiac committed progenitors (**Fig. 4D**). Given that CD82 is a key marker of CM fate specification^43^, these data align with an early role for the lncRNA in CM specification, which is also coherent with the emergence of default alternative fates observed by transcriptomic analyses. Along the same direction, flow cytometry analysis performed at later stages (D20) shows an 80% reduction of differentiated CM, as inferred by the quantification of TNNT2^44^ positive (+) cells (**Fig. 4E** and **Supplementary Fig. 4D**). These findings perfectly match with cellular deconvolution analysis used to interpret our iPSC-derived CM in terms of cell-type composition. Indeed, we found that *pCHARME* ablation causes a reduction of approximately 75% of CM (D20) (**Fig. 4F** and **Supplementary Table 1**), with most of the downregulated DEGs enriched in this population (KL-GSEA, **Supplementary Fig. 4E**). However, as the impact of *pCHARME* ablation in CM was higher than in precursor cells, we asked whether the lncRNA may also play an intrinsic role in differentiated cells. To this end, we used antisense LNA-GapmeRs to deplete *pCHARME* directly in CM (D20). Our findings indicate that 80% reduction of *pCHARME* reduces, although to a lesser extent than in the KO cells, the expression of genes induced with differentiation (i.e *TNNT2*, *MYH7* and *CACNA1C*) (**Fig. 4G**). These findings indicate that *pCHARME* plays a dual regulatory role in the acquisition and maintenance of CM identity by regulating the expression of genes which are crucial for the development and for the physiology of cardiac muscle.

**Fig. 4.**
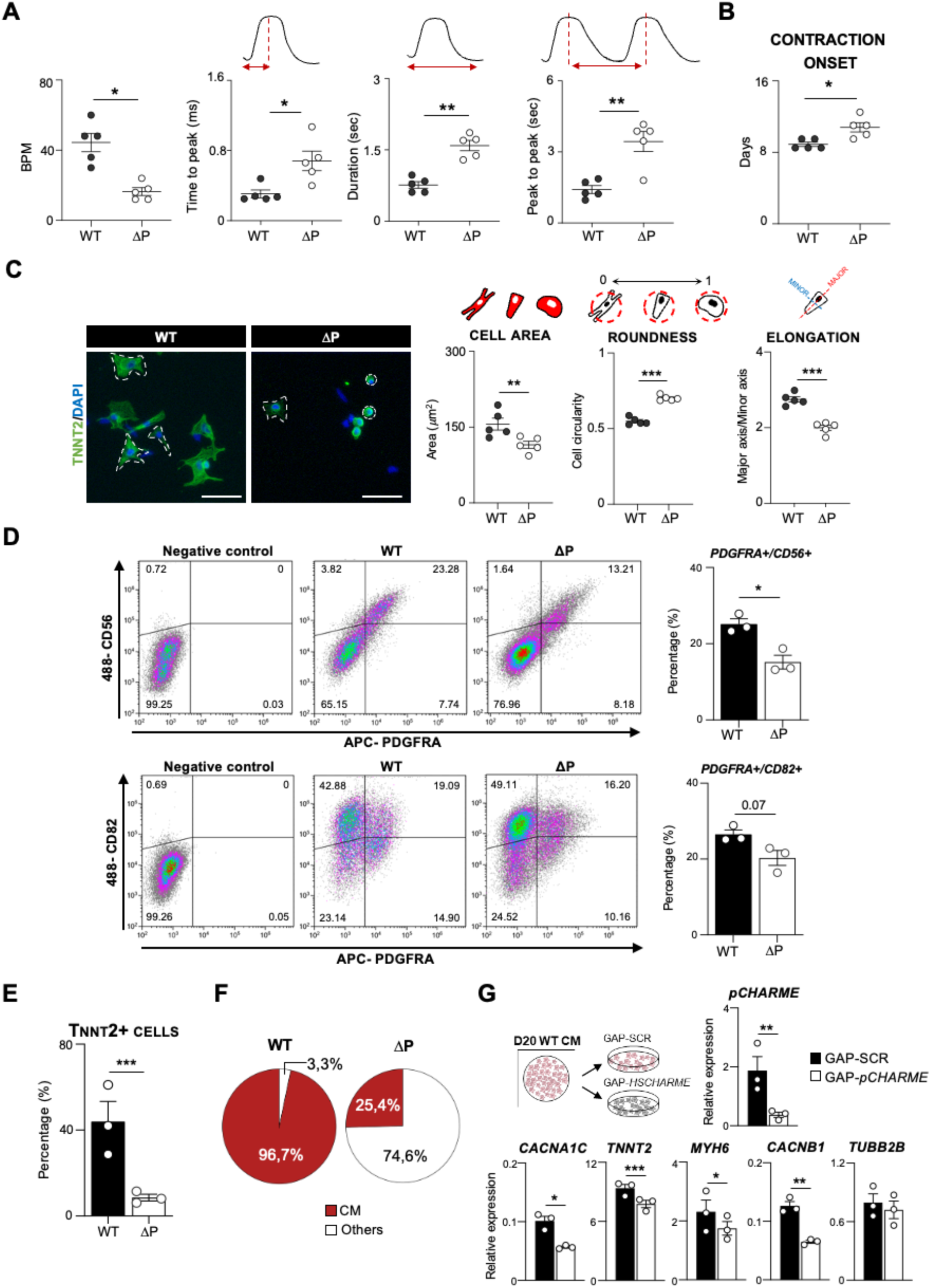
Functional implication of *HSCHARME* in cardiomyogenesis. **A)** Quantitative analysis of the beat rate (expressed as beats per minute=BPM) and contraction parameters in WT and ΔP D20 CM. Representative images of the specific parameter are shown above each graph. Data represent means ± SEM of 5 biological replicates. **B)** Onset (days) of spontaneous contraction of WT and ΔP hiPSC-derived CM. Data represent means ± SEM of 5 biological replicates. **C)** Left panel: Representative images of WT and ΔP hiPSC-derived D20 CM selected for morphological analyses (white dashed outlines) after TNNT2 (green) and DAPI (blue) staining. Right panel: Population measurements of CMs morphological features. A schematic representation of the specific measure quantified is shown above each plot. Black bars represent means ± SEM of 5 biological replicates. **D)** Left panels: Representative flow-cytometry density plot of WT and ΔP hiPSC-CM analyzed at day 10 for cardiac progenitors (PDGFRA/CD56) and cardiac-committed progenitors (PDGFRA/CD82) marker expression. Percentages of each cell population are shown. Right panels: PDGFRA+/CD56+ and PDGFRA+/CD82+ positive cells were set by analysing negative control samples stained with secondary antibody only. Percentages are plotted on the right and represent mean ± SEM of 3 biological replicates. Data information: *p < 0.05; one-sample two-tailed Student’s t-test on LogFC against the null hypothesis of zero (no change). **E)** Flow cytometry quantification of cardiac troponin-T (TNNT2) positive cells percentage in WT and ΔP hiPSC-derived D20 CM. Data represent mean ± SEM of 3 biological replicates. Data information: ***p < 0.001; one-sample two-tailed Student’s t-test on LogFC against the null hypothesis of zero (no change). **F)** Pie-chart displays the estimated proportion of CM (red) and other cell types (white, **Supplementary Table 1**) based on deconvolution analysis of WT and ΔP CM (D20). **G)** RT-qPCR quantification of *pCHARME* and selected mRNA in CM (D20) transfected with control (GAP-SCR) or *pCHARME*-targeting (GAP-*pCHARME*) antisense LNA-GapmeRs. See ‘**Materials and methods**’ for details. Data were normalized to *ATP5O* mRNA and represent relative expression means (2^^-DCt^) ± SEM of 3 biological experiments. Statistical tests were conducted on log₂-transformed fold change (LogFC) values compared to the control sample (GAP-SCR). Data information: *p < 0.05, **p < 0.01, ***p < 0.001; one-sample two-tailed Student’s t-test against the null hypothesis of zero (no change).

### *pCHARME* regulates the alternative splicing of cardiac-specific pre-mRNAs

The high-local distribution of *pCHARME* in the nucleus led us to determine whether specific features are associated with these lncRNA-enriched domains. We first explored the potential for *pCHARME* as a splicing regulator, given the similar sarcomere and contraction defects that splicing factor loss often causes in the heart^45^. In this direction, we combined RNA-FISH targeting *pCHARME* with SC35 immunofluorescence (IF) to determine the proximity of the lncRNA to SC35-containing nuclear speckles^46,47^. Intriguingly, in-depth quantification of three-dimensional signal distribution revealed that *pCHARME* stains in close contact with SC35 domains with 40% of *pCHARME* colocalising with SC35 domains and 60% of SC35 domains containing *pCHARME* (**Fig. 5A-B**).

**Fig. 5.**
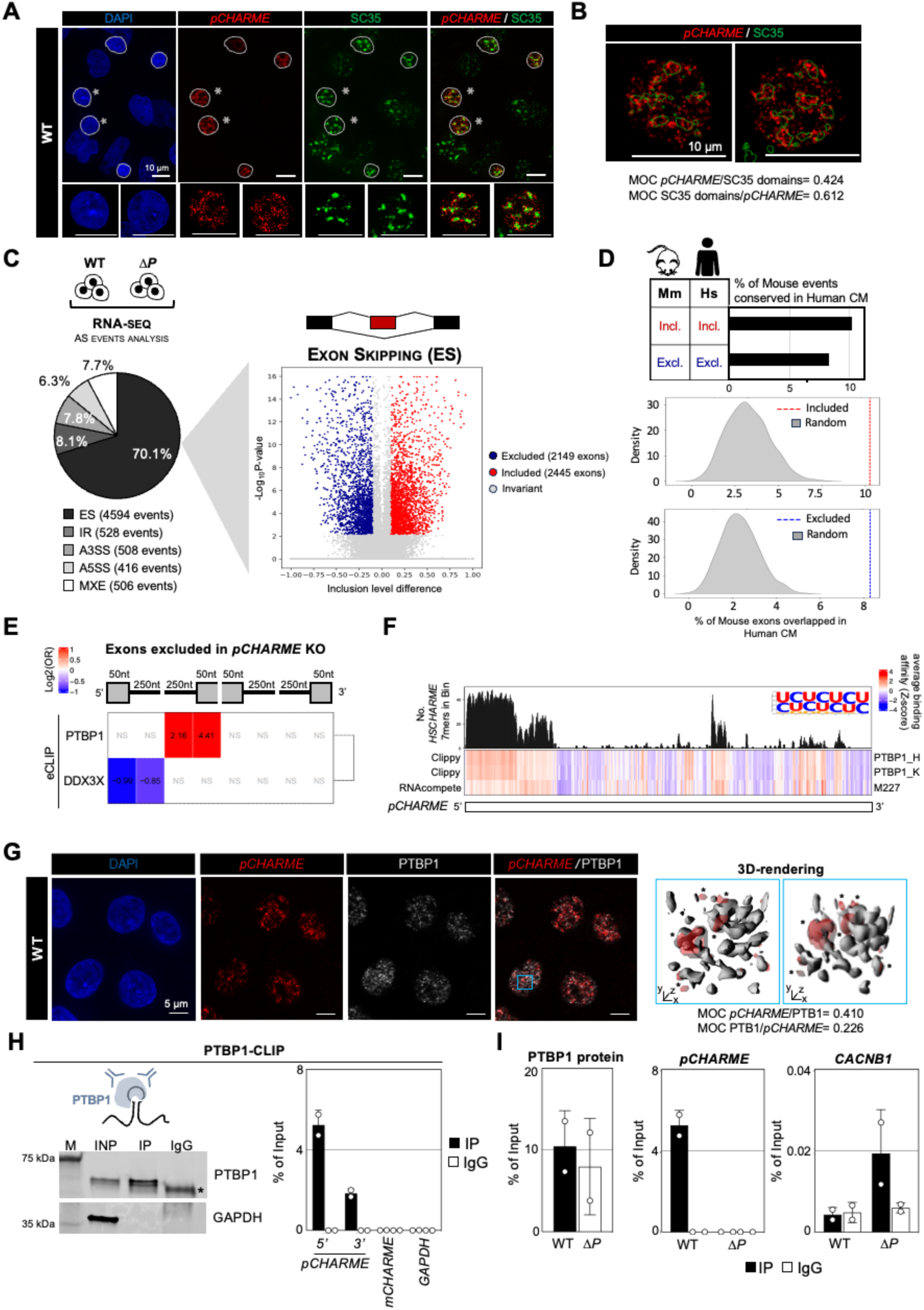
*pCHARME* regulates the alternative splicing of cardiac-specific pre-mRNAs. **A)** Representative maximum 2D projection of the full-size images of confocal caption for RNA-FISH *pCHARME* (red) staining combined with SC35 (green) and DAPI (blue) IF in WT D20 CM. The white lines indicate *pCHARME*-positive nuclei. Digital magnifications of the nuclei highlighted by white asterisks are reported below. **B)** Graphic visualization of *pCHARME*/SC35 domains colocalization signals highlighting the boundary of SC35 domains overlaid with *pCHARME*. Colocalization indexes between *pCHARME* and SC35 domains signals are indicated by Mander’s overlap coefficients (MOC). **C)** Left panel: Pie Chart depicting the portion of significantly altered (FDR<0.05) splicing events detected by rMATS comparing WT vs ΔP D20 RNA-seq samples. All classical splicing alterations were investigated, such as Exon skipping (ES); Intron retention (IR); alternative 3’ splice site (A3SS); alternative 5’ splice site (A5SS) and Mutually exclusive exons (MXE). Right panel: Volcano plot depicts significant (FDR<0.05; Inclusion level variations stronger than 10%) ES events in ΔP D20 CM. X-axis represent exon inclusion ratio while y-axis represent –log10 of P-value. A schematic representation of the investigated event is shown above. **D)** Upper panel: Barplot shows the proportion of exon skipping events identified in murine *pCharme* KO hearts and regulated by *pCHARME* in human CM. Events are stratified by the regulatory outcome: exon inclusion (Incl.) and exon exclusion (Excl.). Lower panel: Density plots represent distribution of conserved events expected by chance (grey distribution). The observed conservation is marked by vertical dashed lines, red for inclusion and blue for exclusion. **E)** Upper panel: Schematic representation of the pre-mRNA regions analyzed in the RBP binding enrichment analysis. Lower panel: Heatmap displaying the log₂(Odds Ratio) of eCLIP binding site (enrichment or depletion) in *pCHARME*-regulated skipping exon (SE) events, compared to unaffected control exons, across the analyzed pre-mRNA regions (from left to right: exonic-upstream, intronic-upstream, intronic-center(5’SS), exonic-center(5’SS), exonic-center(3’SS), intronic-center(3’SS), intronic-downstream and exonic-downstream). Statistically significant enrichment or depletion is indicated by red or blue colors, respectively, while white denotes the absence of statistical significance (FDR > 0.05), also marked by the letters “NS” (Not Significant). Statistical significance was determined using Fisher’s exact test, and p-values were corrected for multiple testing using the Benjamini-Hochberg false discovery rate (FDR) method. **F)** Heatmap displaying the PTBP1 binding affinity for *pCHARME* RNA sequence. The sequence is represented from 5’-end (left) to 3’-end (right) in segments of 50 nucleotides, with a 10-nucleotide sliding step. Bar plot in the top panel displays the total number of occurrences of *pCHARME*-specific 7-mers within each defined segment of the transcript. On the right, the sequence logo generated from the 16 distinct 7-mer types that constitute the *pCHARME*-specific signature is shown. Heatmap below shows the row-scaled (Z-score) PTBP1 binding affinity scores for *pCHARME* segments, based on data from two eCLIP experiments (HepG2 and K562) and one RNAcompete experiment (M227_0.6). Binding affinity scores were calculated as the average PEKA score of all 5-mers within each 50-nucleotide segment for the Clippy-analyzed eCLIP datasets, and as Z-scores for the RNAcompete experiment. **G)** Left panel: Representative confocal images showing RNA-FISH staining for *pCHARME* (red) combined with IF for PTBP1 (white) and DAPI (blue) in WT (D20) CM. Right panel: composite image of *pCHARME* (red) and PTB1 (grey) as volume view of the region depicted with in the blue square. Asterisks point out *pCHARME* regions overlaid with PTBP1. Colocalization indexes between *pCHARME* and PTBP1 signals are indicated by Mander’s overlap coefficients (MOC). **H)** Left panel: Western blot analysis of PTBP1 protein from PTBP1-CLIP assay in WT D20 CM. GAPDH protein serves as a loading control. Input (Inp) samples represent 10% of the total protein extracts. Right panel: RT-qPCR quantification of *pCHARME* (amplified at both 5′ and 3′ intron-1 ends) and *mCHARME* recovery in PTBP1 IP and IgG samples. *GAPDH* RNA serves as negative control. Values are expressed as percentage of input and represent mean ± SD of 2 biological replicates. I) Left panel: PTBP1 protein recovery in WT and ΔP D20 CM. Quantification of the chemiluminescent signal was performed with the ImageJ tool. Values are expressed as percentage of input and represent mean ± SD of 2 biological replicates. Middel panel: RT-qPCR quantification of *pCHARME* (amplified at 5′ intron-1 end) recovery in PTBP1 IP and IgG samples from WT and ΔP D20 CM. Values are expressed as percentage of input and represent mean ± SD of 2 biological replicates. Right panel: RT-qPCR quantification of *CACNB1* RNA recovery in PTBP1 IP and IgG samples from WT and ΔP D20 CM. Values are expressed as percentage of input and represent mean ± SD of 2 biological replicates.

The observed RNA-protein proximity led us to test a possible involvement of *pCHARME* in splicing regulation. By large-scale analysis of splicing patterns from our RNA-seq datasets, we identified a total of 9615 expressed isoforms in both WT and ΔP cardiomyocytes (see **Materials and Methods**). We found that the abundance of a consistent fraction of these isoforms (16.3%; corresponding to 1877 transcript isoforms) was significantly altered upon *pCHARME* ablation and functionally linked to cardiomyogenic ontologies related to cardiac muscle cell development and heart contraction (**Supplementary Fig. 5A**). On this preliminary evidence, we used rMATS^48^ for alternative splicing (AS) analysis of our RNA-seq datasets, including exon-skipping (ES), intron retention (IR), alternative 3’ splice site (A3SS), alternative 5’ splice site (A5SS), and mutually excluded exons (MXE) and events. Also in this case, we observed a strong impact of *HSCHARME* ablation, with a total of 6.552 aberrant pre-mRNA splicing events (FDR<0.05, Absolute Inclusion level>0.1) (**Fig. 5C**, **Supplementary Fig. 5B** and **Supplementary Table 3**). Splicing alterations were found in each of the analysed classes, with the ES events accounting the 70.1% of all the significant alterations (**Fig. 5C**, right panel). Among the most interesting examples of cardiac genes, we found *CACNB1*, a gene implicated in voltage-dependent calcium release^49^. In KO CM (ΔP and PA), we found that the skipping of exon 7 of *CACNB1* causes a shift from the EX6-EX7-EX8 isoform into the shorter EX6-EX8 one (**Supplementary Fig. 5C-D**). Another example is *MYL6*, a gene encoding for a hexameric ATPase cellular motor protein expressed in muscle and non-muscle tissues^50^. *MYL6* gene produces two mRNA variants which differ in exon 6^51^ (**Supplementary Fig. 5E**). We found that *HSCHARME* KO (ΔP and PA) leads to the overabundance of the longest EX5-EX6-EX7 isoform (**Supplementary Fig. 5E-F**), which is described in the literature as the major non-muscle transcript. Therefore, the *HSCHARME*-mediated regulation of MYL6 splicing facilitates the production of a structural myosin variant optimized for CM contraction.

To evaluate whether *pCHARME* regulation of alternative splicing is also evolutionarily conserved between human and mouse CM, we performed AS analysis of RNA-seq datasets from WT and *Charme* KO post-natal hearts (GSE200878^22^). We found that, *in vivo*, the lack of *Charme* results in 327 exon exclusion and 265 exon inclusion events (**Supplementary Fig. 5G** and **Supplementary Table 3**). We mapped the coordinates of these differentially spliced murine exons to the human genome and searched for overlaps with *pCHARME*-regulated exons. Focusing on concordant events, we observed conservation for 10.2% of inclusion events and 8.3% of exclusion events (**Fig. 5D**, upper panel). Notably, these proportions were significantly higher than expected by chance (**Fig. 5D**, lower panel), underscoring the conserved regulatory role for the lncRNA. Among the conserved splicing targets, we identified key cardiac genes, including *TNNT2*^44^, *MFF*^52^, *SYNC*^53^, *SORBS2*^54^, *CREM*^55^ and *RBFOX2*^56^, which are known to play crucial roles in heart function (**Supplementary Table 3)**. Overall, these results indicate *pCHARME* as a novel, evolutionarily conserved regulator of CM splicing.

### *pCHARME* modulates alternative splicing by direct interaction with PTBP1

To gain mechanistic insights into the splicing alterations observed in the absence of *pCHARME*, we leveraged data from 223 eCLIP experiments covering 150 RNA-binding proteins (RBP) across cell lines (K562 and HepG2)^57^ that do not encode the lncRNA. By focusing on ES, which represent the most altered and conserved *pCHARME*-regulated splicing events, and comparing them to a control set of similarly expressed but unregulated junctions (**Supplementary Fig. 5H**), we identified the splicing repressor PTBP1 as significantly enriched at the 5′ junctions of the excluded exons (Fisher’s exact test, FDR < 0.05, log2[Odds Ratio] > 0) (**Fig. 5E** and **Supplementary Table 4**). Among them we found several known PTBP1 splicing targets, such as *TPM1*, *ACTN1*, *FHOD3*, and *TNNT2*^58^, expressed in CM and displaying exon exclusion in *pCHARME* KO (**Supplementary Table 4**).

Overall, these data prioritise PTBP1 as a regulator of the *pCHARME*-dependent ES events and suggest a possible interplay between the two in CM. To deepen this evidence with a focus into the physical engagement of *pCHARME*, we analyzed its nucleotide composition (7-mers) compared to the set of the 1987 lncRNAs expressed in CM (**Supplementary Fig. 5I**). We found that *pCHARME* possesses a unique UC-rich sequence signature, primarily driven by an extended simple repeat (75-2955 nt) in the 5’ region of intron 1 (**Supplementary Table 4**). RNA secondary structure predictions using the RNAfold software^59^ suggested these UC-rich stretches as unstructured and likely accessible to RBP (**Supplementary Fig. 5J**). On this prediction, we employed four independent approaches to identify RBP with high affinity for the UC-rich sequence, and specifically (i) RBP affinity scores derived from crosslinking-induced truncations from eCLIP datasets, (ii) *in vitro* RNAcompete experiments assessing RBP binding preferences for short nucleotide sequences^60^, (iii) in silico predictions, and (iv) motif analysis using CatRAPID software^61^. The results showed only two RBP (PTBP1 and PCBP2; **Supplementary Fig. 5K** and **Supplementary Table 4)** as the most likely interactors of the *pCHARME* UC-rich signature. Notably, PTBP1, previously found as a *pCharme* interactor in murine myocytes^21^, showed binding propensity scores closely mirroring the abundance of the 5’ UC-rich sequences (**Fig. 5F**).

In support of the PTBP1/*pCHARME* interaction, we performed PTBP1-IF assay combined with *pCHARME* RNA/FISH in CM, which revealed that approximately 40% of the lncRNA signal overlapped with PTBP1 (**Fig. 5G**). Additionally, we tested for a direct interaction between PTBP1 and the lncRNA by using UV cross-linking immunoprecipitation (CLIP) experiments in CM. Western blot analysis confirmed successful recovery of PTBP1 after immunoprecipitation (**Fig. 5H**, left panel). Subsequent RT-qPCR analysis of the retrieved RNA using isoform-specific primers confirmed a robust and specific interaction between PTBP1 and the 5′ region of *pCHARME* intron 1 (**Fig. 5H**, right panel), which aligns with the presence of the UC-rich motifs and the highest PTBP1 binding propensity.

Together with the observation that PTBP1 binds *pCHARME* splicing-regulated targets in the absence of the lncRNA, this PTBP1/*pCHARME* interaction in CM suggests a role for the lncRNA as a decoy for PTBP1. To test this hypothesis, we examined the potential binding of PTBP1 to the *pCHARME* splicing target *CACNB1*, whose exon 7 was excluded upon the lncRNA ablation (**Supplementary Fig. 5C-D)**. Supporting the decoy mechanism, RT-qPCR analysis from PTBP1-CLIP experiments performed in WT and KO conditions shows the specific binding of PTBP1 at the level of *CACNB1* exon 7 in KO CM, while no enrichment was found in WT cells (**Fig. 5I**). As PTBP1 was recently shown to negatively regulate CM specification^62^, the suggested role of *pCHARME* as a PTBP1 decoy could support a mechanism ensuring the acquisition of splicing patterns that are specific for CM.

### *HSCHARME* dosage impacts the expression of disease-linked genes

The influence of *pCHARME* in CM differentiation encouraged further investigation into the role of the human transcript in pathology. GSEA analyses performed on *HSCHARME* DEGs (*Δ*P *vs* WT) evidenced a substantial enrichment of *HSCHARME* targets among genes dysregulated in HCM (**Fig. 6A**, left panel) and DCM (**Fig. 6A**, right panel) cardiomyopathies. Further confirmation was obtained by KEGG pathways enrichment analysis performed on the same subsets, showing a significant clustering of *HSCHARME* targets into the HCM (adjusted p-value 1.51E^-10^) and the DCM (adjusted p-value 3.86E^-9^) categories (**Fig. 6B**). These findings strongly suggest that the human gene may play a role in the development or the homeostasis of these cardiac pathological states. To strengthen this hypothesis, we performed DE analyses of available RNA-seq datasets of HCM and DCM cardiomyopathies and Healthy human cohorts (GSE130036^63^; GSE116250^64^). Interestingly, we found aberrant expression of the *HSCHARME* locus with a significant up-regulation of the *pCHARME* isoform in both HCM and DCM diseased hearts (**Fig. 6C**).

**Fig. 6.**
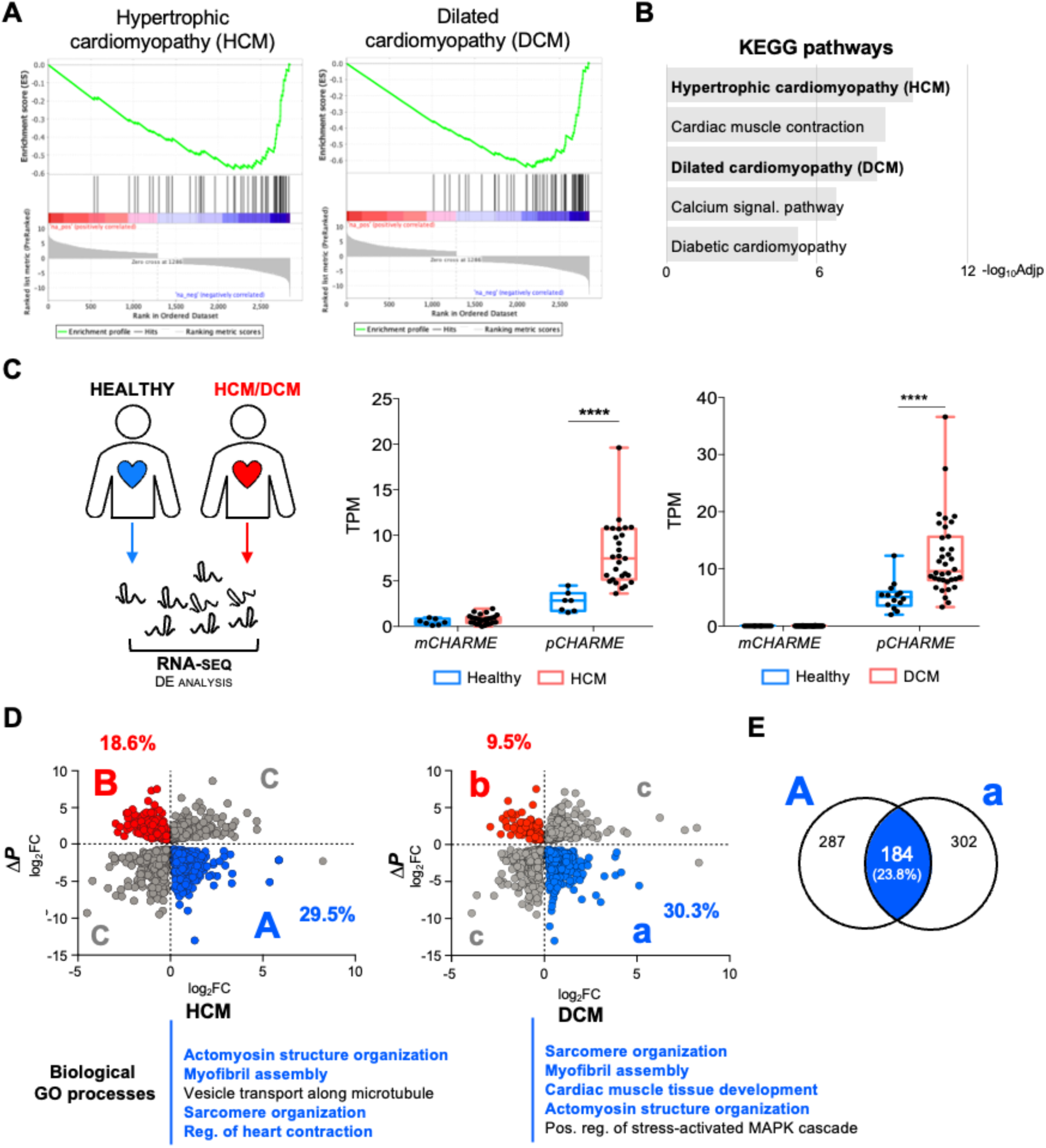
Identification of *pCHARME* targets in hypertrophic and dilated cardiomyopathies. **A)** GSEA plot showing the enrichment of the Human Phenotype Ontology terms “Hypertrophic Cardiomyopathy” and “Dilated Cardiomyopathy” that resulted strongly down-regulated in the analysis of ΔP vs WT D20 CM. **B)** KEGG pathways enrichment analysis performed with EnrichR on significant DEGs from ΔP vs WT D20 CM. Bars indicate the top categories of biological processes in decreasing order of –log10 adjusted p-value (–log10Adjp). All the represented categories show an Adjp value <0.05. **C)** Quantification of *mCHARME* and *pCHARME* expression from RNA-seq analysis performed in healthy (CTRL) vs patients of hypertrophic (left panel) and dilated (right panel) cardiomyopathies. Data information: *p < 0.05; ***p < 0.01; ****p<0.0001; FDR **D)** Scatterplots representing the log2FC comparison between DEGs in ΔP and HCM (left panel) and DCM (right panel), showing up- and down-regulated genes with opposite trends in the conditions. Percentage of de-regulated genes are reported as well as Biological Process GO categories in which they cluster. All reported categories show an Adjp <0.05. **E)** Venn diagrams depicting the intersection between the “A” (ΔP vs WT down-regulated DEGs crossed with Healthy vs HCM up-regulated DEGs) and “a” (ΔP vs WT down-regulated DEGs crossed with Healthy vs DCM up-regulated DEGs) categories.

To more directly pinpoint the genes whose aberrant expression in pathology is influenced by *pCHARME* alteration, we then used a guilt-by-association approach and compared the lists of *HSCHARME* targets (**Supplementary Table 3**) to HCM and DCM DEGs (**Supplementary Table 5**). We observed more than 1500 significant DEGs for each comparison, n=1598 for HCM and n=1603 for DCM (**Fig. 6D**). Among them, the expression of 29.5% disease-associated DEGs (category “A”, **Fig. 6D**, left panel) was correlated with *pCHARME*, being up-regulated in HCM and down-regulated in the lncRNA knockout cells. Intriguingly, these DEGs clustered into categories of interest for heart functionality, such as actomyosin structure organization, myofibril assembly, sarcomere organization and regulation of heart contraction. Notably, the “A” category also contains the *CACNB1* transcript whose splicing pattern was significantly altered upon *HSCHARME* ablation (**Supplementary Fig. 5C-D**). Similarly, the comparison with DCM revealed that 30.3% of DEGs (category “a”, **Fig. 6D**, right panel) were up-regulated in the DCM cohorts and significantly down-regulated upon *HSCHARME* ablation. Again, these genes clustered into categories functionally related to heart development, such as sarcomere organization, myofibril assembly and cardiac muscle tissue development. These results are in line with recent evidence demonstrating the central role of CM and the contractile apparatus in DCM^65^ and highlight the potential involvement of *pCHARME* in disease.

On the other hand, the intersection between genes up-regulated upon *HSCHARME* ablation and down-regulated in pathologies was limited to a few cases with little interest in CM development (categories “B” and “b”, **Fig. 6D**). The remaining DEGs (categories “C” and “c”, **Fig. 6D**) display expression trends that are opposite to those of *pCHARME*. Consequently, their changes in pathology cannot be attributed to *pCHARME* upregulation but are likely influenced by other context- or pathology-specific effects. The use of the same compound in the treatment of cardiomyopathy can constitute a novel opportunity for therapeutic applications. In this direction, the comparison of DEGs included in the categories “A” and “a” led to the discovery of a common subset of cardiac homeostasis and contraction genes (n=184; **Fig. 6E**). Notably, these transcripts, whose expression in CM was activated by *HSCHARME* and, coherently with its pathological upregulation, aberrantly overexpressed in both HCM and DCM hearts. This subset represents a group of candidates for developing therapeutic strategies. With the aim to test their responsiveness to the modulation of *pCHARME* levels, we established an inducible hiPSCs CRISPR-based system (RHE) acting as a dynamic rheostat to either silence or induce *HSCHARME* within the same genetic background. CRISPR-cas9 mediated homologous recombination was achieved by designing a single sgRNA (**Supplementary Table 2**) to add the doxycycline (DOXY)-responsive element (TRE promoter) upstream of the *HSCHARME* locus (**Supplementary Fig. 6A**). Assessment of *pCHARME* expression by RT-qPCR and RNA-FISH analyses confirmed the success of the editing strategy, as the lncRNA levels were comparable to knockout cells in the untreated CM and increased significantly upon doxycycline addition (-/+ DOXY; **Fig. 7A**). To gain functional validation on the impact of *pCHARME* dosage in RHE-CM, we then quantified the expression of several lncRNA targets identified by our RNA-seq analysis (**Fig. 3C** and **Supplementary Fig. 3G**). In line with our loss-of-function analyses, we focused on genes critical for CM structure (i.e. *MYH7*, *TNNT2*) and functionality (*CACNA1C*, *CACNB1*). RT-qPCR analyses revealed that their expression was consistently modulated by *pCHARME* levels, decreasing in -DOXY conditions while triggered upon +DOXY treatments (**Supplementary Fig. 6B**). This effect was highly specific as demonstrated by *TBX5* which levels were found unaltered, in line with its role as an upstream regulator of the lncRNA expression (**Supplementary Fig. 6B**). Notably, these *pCHARME*-responsive genes (i.e. *MYH7*, *TNNT2* and *CACNA1C*) are recognized as known cardiomyopathy-causative loci^66,67^ with essential roles in maintaining CM functionality and homeostasis. Consequently, the ability to regulate their expression *via* modulation of *pCHARME* levels offers a promising avenue to ameliorate the pathological state of patients across various genetic backgrounds. Finally, we extended the analysis to our priority list of targets common to “A” and “a”, therefore consistently de-regulated in both DCM and HCM (**Fig. 6E**). Intriguingly, we found that genes such as *PPFIA4*, *NPPB, CAMK2B* and the cardiomyopathy-linked *MYO18B*^68^, displayed significant and coherent responses to *pCHARME* modulation (**Fig. 7B**). Finally, we also analysed the responsiveness of these disease-linked genes to the transient knockdown of *pCHARME* in differentiated CM and found the consistent decrease of all of them (**Fig. 7C**). Overall, the ability of *pCHARME* to influence gene expression programs, including the regulation of cardiac genes linked to disease, marks an advancement in the lncRNA field and suggests its potential relevance for future studies exploring diagnostic or therapeutic strategies.

**Fig. 7.**
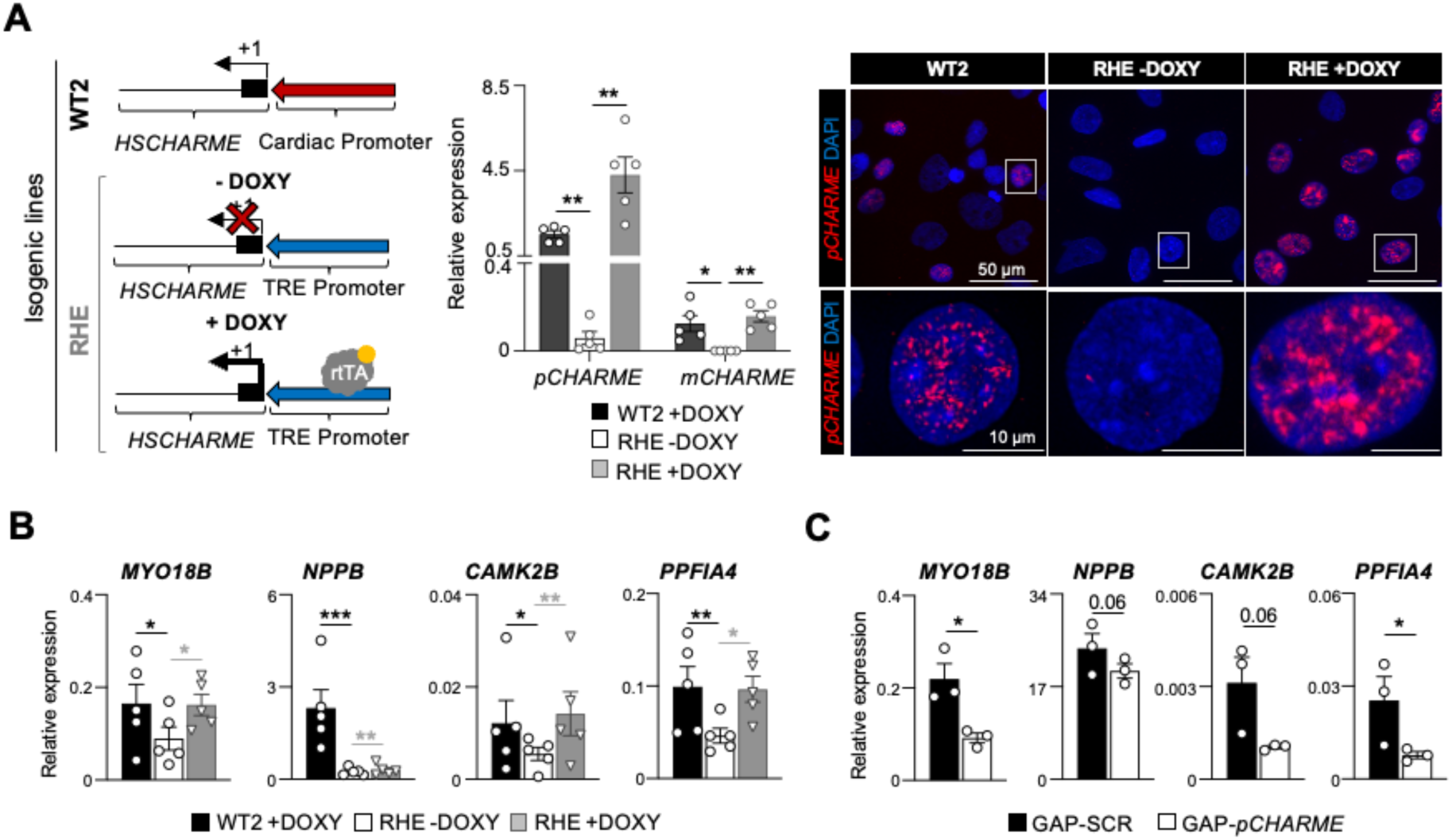
*pCHARME* dosage impacts the expression of disease-linked genes. **A)** Left panel: Schematic representation of the CRISPR/Cas9 genome editing strategies used to obtain the RHE-hiPSCs. Middle panel: RT-qPCR quantification of *pCHARME* and *mCHARME* expression in WT2, RHE (-DOXY) and RHE (+DOXY) hiPSC-derived CM. Data were normalized to *ATP5O* mRNA and represent relative expression means (2^^-DCt^) ± SEM of 5 biological replicates. Right panel: Representative maximum 2D projection of the full-size image of confocal caption for *pCHARME* RNA-FISH (red) and DAPI (blue) staining in WT2, RHE (-DOXY) and RHE (+DOXY) CM (D20). Digital magnifications of the nuclei highlighted by white squares are reported below. **B)** RT-qPCR quantification of pathology-linked transcripts in WT2, RHE (-DOXY) and RHE (+DOXY) hiPSC-derived CM (D20). Data were normalized to *ATP5O* mRNA and represent relative expression means (2^^-DCt^) ± SEM of 5 biological replicates. Statistical tests were conducted on log₂-transformed fold change (LogFC) values compared to the control sample (RHE -DOXY). **C)** RT-qPCR quantification of pathology-linked transcripts in WT CM (D20) transfected with control (GAP-SCR) or *pCHARME*-targeting (GAP-*pCHARME*) antisense LNA GapmeRs. See ‘**Materials and methods**’ for details. Data were normalized to *ATP5O* mRNA and represent relative expression means (2^^-DCt^) ± SEM of 3 biological experiments. Statistical tests were conducted on log₂-transformed fold change (LogFC) values compared to the control sample (GAP-SCR). Data information: *p < 0.05; ***p < 0.01; ****p<0.0001; one-sample two-tailed Student’s t-test against the null hypothesis of zero (no change).

## DISCUSSION

Understanding the molecular causes of cardiac pathologies represents a priority for human health. However, the inability of the heart to regenerate and its mixed cellularity add complexity to medical management, making cardiomyopathies the leading cause of death in developed countries^69,70^. In this context, lncRNAs are emerging as promising candidates for fine-tuning CM-specific gene expression programs due to their precise spatiotemporal regulation^71^. New biopharmaceutical companies are starting to show the potential of lncRNA-based platforms in enhancing precision medicine strategies for a wide range of illnesses influenced by lncRNA levels, including cardiovascular diseases. This is the case of the HAYA’s therapeutic candidate HTX-001, an antisense oligonucleotide targeting the heart-specific lncRNA Wisper^72^.

Here, we provide evidence of the importance of *HSCHARME* in the pathophysiology of the heart and set the stage for innovative approaches of lncRNA-targeting precision therapies for the cell-specific treatment of human cardiomyopathies (**Fig. 8**). Indeed, the conservation across species and the high specificity of *HSCHARME* expression for CM make this lncRNA an excellent candidate for pioneering therapeutical treatments that could minimise off-target effects on other cardiac cell types. Currently, several examples of lncRNA playing a role in human CM homeostasis have been identified^5,6,10,73^. However, these studies rely on *in vitro* systems that induce differentiation exclusively towards cardiomyocytes. Therefore, it is unclear whether the role of these lncRNAs in cardiac processes is truly cardiomyocyte-specific or if they could also have roles in other cardiac cell subtypes, such as cardiac fibroblast or endothelial cells.

**Fig. 8.**
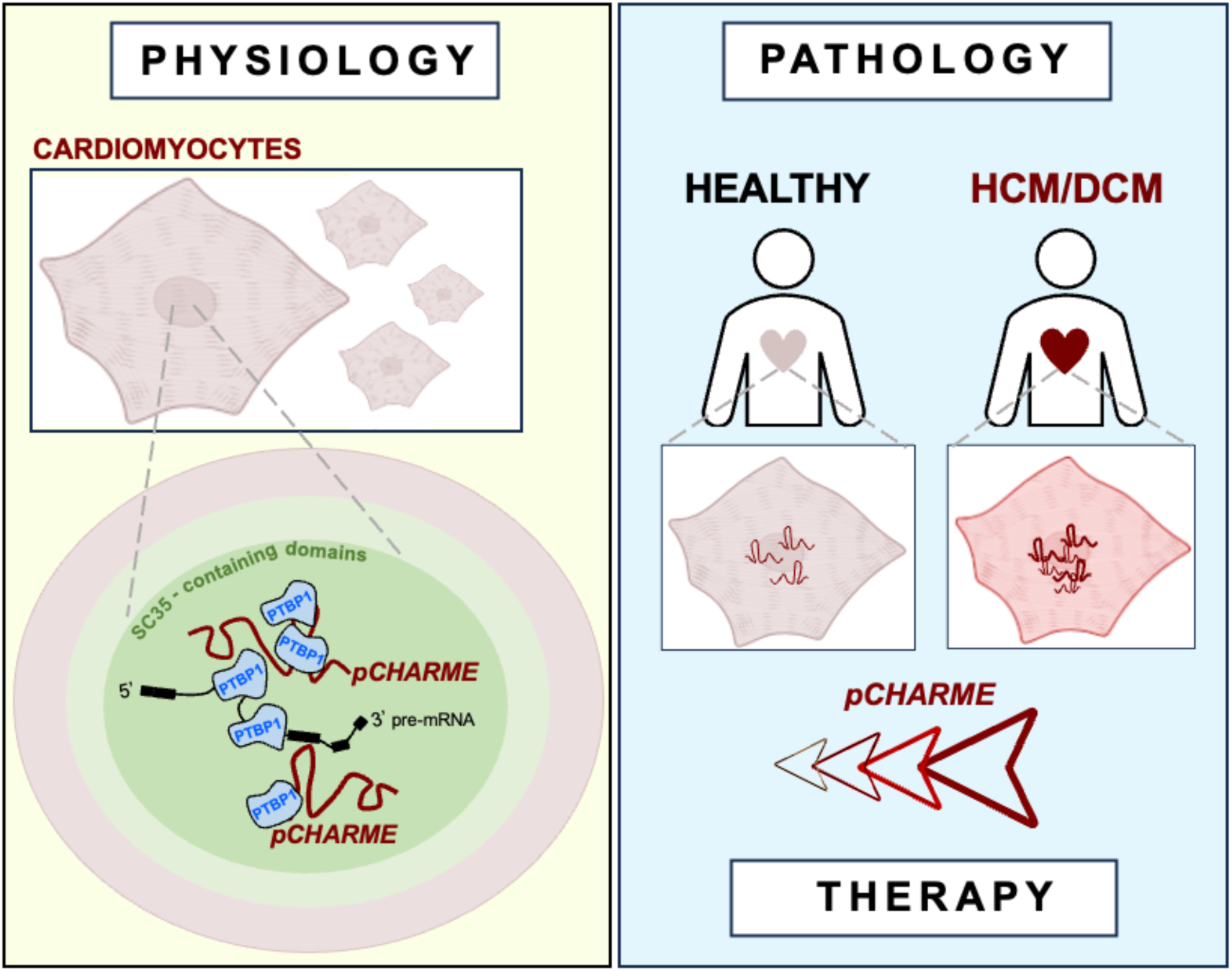
Overview of *pCHARME* role in cardiomyocyte physiology and disease. Under physiological conditions (Physiology) *pCHARME* localises within S35-domains and acts as a decoy for the splicing repressor PTBP1, thereby positively regulating the expression of genes involved in the differentiation of human cardiomyocytes. In DCM and HCM cardiomyopathy hearts (Pathology), *pCHARME* is aberrantly upregulated, and its target genes -many of which are associated with the diseases-are also dysregulated. This scenario envisages novel therapeutic approaches (Therapy) aimed at regulating disease-linked genes through *pCHARME* modulation.

In our study, publicly available single-cell datasets allowed us to uncover the specific expression of *HSCHARME* in human CM, providing a solid basis to investigate its role more accurately and specifically within the cardiac context. We found that the human syntenic *pCHARME* transcript shares a high level of conservation with its murine counterpart and maintains peculiar features that make the lncRNA functional, such as the retention of intron 1. Mechanistically, we provide evidence of the direct interaction between *pCHARME* and the splicing suppressor PTBP1 and show their functional interplay in regulating a subset of the lncRNA targets at the splicing level. Given the strong binding affinity between PTBP1 and *pCHARME*, we propose a model in which the lncRNA acts as a decoy for PTBP1, sequestering the splicing repressor from pre-mRNA targets, thereby promoting the inclusion of specific exons and ensuring the acquisition of CM-specific splicing patterns. This model aligns with the recently described role of PTBP1 as a negative regulator of CM specification^62^. As such, the interaction between *pCHARME* and PTBP1 underscores the importance of this axis in maintaining proper cardiac splicing programs and highlights its potential relevance in cardiac pathophysiology. Recently, nuclear speckles have been described as storage sites for transcripts with retained introns^74^. These unspliced RNAs retain certain structural features, such as a high GC content and a high density of protein binding motifs^75^ that perfectly fit with *pCHARME* and strengthen our proposed model of speckle-associated lncRNA.

As the role of nuclear speckles in regulating cellular specification is still only partially understood, the muscle-specific restriction of *pCHARME* can add specificity to their implication in cardiac-specific gene expression programs. Remarkably, as for *HSCHARME* (this paper), alterations in SC35 expression were linked to dilated cardiomyopathy^76^, thereby narrowing a possible line of intervention for cardiac disorders. To note, the role of *pCHARME* in regulating structural genes, such as different types of myosins and troponins, is also in line with a broader role for the human lncRNA in the functionality of muscle cells, either cardiac or striated, as previously characterized in mice^20,21^. A newly identified shorter isoform of *pCHARME* was identified in human myocytes as a regulator of *MYBPC2* expression (lncFAM^23^). However, the fact that *MYBPC2* is neither expressed nor influenced by *HSCHARME* in CM (data not shown) would suggest a different role for the two transcripts in the heart.

We also provide evidence that modulating *pCHARME* expression in differentiated CM can influence the levels of several disease-linked genes involved in CM physiological processes, such as calcium handling, heart contraction and CM development. Given its role as a positive regulator of CM differentiation, the modulation of *pCHARME* expression levels might be beneficial for guiding CM de-differentiation into a progenitor-like state, which is crucial for their replenishment after injury and, ultimately, for the effective regeneration of the cardiac muscle *in vivo*^77–79^. Lastly, the proven association between *pCHARME* dysregulation and cardiac pathologies, indicates the necessity of including the lncRNA in genetic testing, particularly for cardiomyopathies with no clear genetic evidence. Alternatively, assessment of *HSCHARME* expression levels could assist in identifying differences in outcome prediction when stratifying cohorts of cardiomyopathy patients who have tested positive for genetic markers, based on phenotype manifestation.

## MATERIALS AND METHODS

### Cell culture conditions and transfection

Human Episomal iPSCc (WT from ^80^ and WT2 are the WTSIi004-A cell line from EBiSC; https://ebisc.org/WTSIi004-A) were maintained in Essential 8 media (Life Technologies) in plates coated with Geltrex LDEV-free reduced growth factor basement membrane matrix (Life Technologies). Differentiation into cardiomyocytes was performed as described in ^32^. Specifically, cells were grown as monolayers in RPMI 1640 Medium (Life Technologies) supplemented with B27 Supplement without Insulin (Life Technologies) for 10 days and then switched to RPMI 1640 Medium (Life Technologies) supplemented with B27 Supplement with Insulin (Life Technologies). The induction towards the cardiac lineage was performed by upregulating canonical Wnt signalling (adding CHIR-99021; Selleck) in the early stages of differentiation and by its subsequent inhibition after 3 days (adding IWR-1; Sigma). Genome editing analyses on hiPSCs were performed by co-transfecting the PX333 plasmid, harbouring the cloned sgRNAs (designed with https://chopchop.cbu.uib.no/; sequences available upon request) and the CAS9 sequence, together with the HR 110PA-1 donor vector (System Biosciences) in which two homology arm sequences were cloned. Cloning was performed employing In-fusion® kit (Takara), according to the manufacturer’s protocol. Cloning with this approach requires adding 15 nt-long tails to the oligonucleotides to create a homology region among insert and plasmid, allowing recombination into bacteria cells (E.Coli STBL3®). The transfection was performed using the Neon Transfection System (Life Technologies). After 48 hours, cells were selected using 1 μg/ml puromycin to identify heterozygous and homozygous *HSCHARME* KO or *HSCHARME* RHE clones. For RNAi, WT hiPSC-derived CM were dissociated into single cells and transfected with either control (GAP-SCR) or *HSCHARME* (GAP-*pCHARME*) antisense LNA GapmeRs (sequences available upon request) using TransIT-X2® Dynamic Delivery System (Mirus). Total RNA was collected 48 hr after transfection.

### RNA extraction and RT-qPCR analysis

Total RNA from cultured cells and tissues was isolated using TRI Reagent (Zymo Research), extracted with Direct-zol^TM^ RNA MiniPrep (Zymo Research), treated with DNase (Zymo Research), retrotranscribed using PrimeScript Reagent Kit (Takara) and amplified by RT-qPCR using PowerUp SYBR-Green MasterMix (Life Technologies), as described in ^21^ (see **Supplementary Table 6**). Data are represented as relative expression (2^^-DCt^). Nucleus/Cytoplasm fractionation of hiPSC-derived CM was performed using the PARIS kit (Life Technologies), following the manufacturer’s specifications.

### RNA-Sequencing Analysis

Total RNA was collected from hiPSCs-derived CM (day 10 and day 20) obtained from three independent biological replicates. Illumina Stranded mRNA Prep was used to prepare cDNA libraries. RNA-Seq was performed on an Illumina Novaseq 6000 Sequencing system at the Genomic Facility of the Center for Human Technologies of IIT.

Q30-filtered raw reads were obtained from Illumina BaseSpace Reads and aligned to GRCh38 assembly using STAR aligner software^81^. Gene loci fragment quantification was performed on Ensemble (release 87) gene annotation gtf using STAR –quantMode GeneCounts parameter. The gtf file was edited adding the *HSCHARME* genomic coordinates (**Supplementary Table 3**). Read counts of “reverse” configuration files were combined into a count matrix file, that was given as input to DESeq2 R package^82^ for normalization and differential expression analysis, after removing genes with less than 10 counts in at least two samples. Adjusted p-value cutoff for selecting significant differentially expressed genes was set to 0.05 unless otherwise specified. Heatmap of differentially expressed genes was generated using pheatmap R package^83^ from normalized scaled data.

Volcano plots were generated using *Enhanced Volcano* R package (https://bioconductor.org/packages/devel/bioc/vignettes/EnhancedVolcano/inst/doc/Enhanced Volcano.html). Gene Ontology enrichment analyses were performed with the EnrichR online tool (https://maayanlab.cloud/Enrichr/^84–86)^ using standard settings. Gene Set Enrichment Analysis was performed with the GSEA tool from Broad Institute^87,88^ running a pre-ranked analysis with statistically significant genes ordered by log2 fold change (KO *vs* WT D20). Gene Ontology was run using the complete GO annotation file (c5.all.v2023.2.Hs.symbols.gmt). Human Phenotype Ontology (HPO) was run using the HPO annotation file (c5.hpo.v2023.2.Hs.symbols.gmt). Figures were prepared with R and Prism 7.0 software and edited with Adobe Illustrator 2024.

### RNA Isoforms Analysis

Reads derived from the WT RNA-seq datasets were used as input for the Salmon tool^89^ to identify and quantify the *HSCHARME* isoforms. A transcriptome file was generated using the human genome FASTA file and the GTF file containing *HSCHARME* coordinates. Transcripts per million (TPM) output values were used for quantification and statistical analysis. Plot for quantifying the *HSCHARME* isoforms was generated with the *ggtranscript* R package (https://github.com/dzhang32/ggtranscript), using the genomic coordinates provided within the GTF annotation file derived from Salmon alignment. Figures were edited with Adobe Illustrator 2024. For large-scale isoform analysis from WT and ΔP datasets, transcript quantification was performed with the Salmon tool using standard parameters and FASTQ files as input. Transcripts per million (TPM) counts were used to quantify the expression of each transcript isoform. Differential gene expression analysis was performed with edgeR^90^, using a quasi-likelihood negative binomial generalized log-linear model (glmQLFit function) and applying a false discovery rate (FDR) cutoff of 0.05 to identify statistically significant genes.

### Single Cell RNA-seq deconvolution analysis

Single-cell RNA sequencing (scRNA-seq) data of hiPSC-derived CM (14 and 45 days of differentiation) were obtained from the Synapse database (ID: syn7818379). FASTQ files were extracted from the provided BAM files using CellRanger BamtoFastq (v1.4) and aligned to the GRCh38 reference genome using Ensembl 99 gene annotation, supplemented with a custom annotation for the long non-coding RNA *HSCHARME*. Cells were filtered to retain high-quality single-cell transcriptomes, defined by having more than 200 and fewer than 5,000 detected genes and less than 20% mitochondrial gene content. The upper threshold for mitochondrial gene content was extended beyond standard filters to retain metabolically active cell types such as CM. Reanalysis was performed with Seurat v5.2.1 in R^91^ the data have been first normalized using NormalizeData function with “LogNormalize” method and then integrated via Seurat’s Canonical Correlation Analysis (CCA) standard workflow.

After integration, initial unsupervised clustering was performed using the Seurat pipeline; Principal component analysis (PCA) was applied on the 3000 most variable genes identified using the VST method, and the top 15 principal components were used to construct a shared nearest neighbour (SNN) graph. Clusters were identified using the Louvain algorithm with a resolution parameter of 0.5. Cell population identities were assigned using a knowledge-based approach by evaluating canonical marker gene expression across clusters. CM were identified based on the expression of known cardiac muscle genes^92^. Neuronal populations were annotated using gene sets associated with the Gene Ontology (GO) terms for postsynaptic density (GO:0014069) and axon part (GO:0033267). Endothelial populations were annotated based on expression of established endothelial markers^62,93,94^. Differential gene expression analysis was performed using the RunPresto function from the Presto package v1.0 in R, applying the Wilcoxon rank-sum test. Each cell population was compared against all remaining cells in a one-vs-all manner. Genes with log2 fold change > 1 and adjusted p-value < 0.05 were considered significantly upregulated. Identified marker genes for each cluster were subjected to functional enrichment analysis using the WebGestaltR R package v0.4.6. The reference gene background used for enrichment consisted of all genes detected in the scRNA-seq experiment after quality control and filtering.

Deconvolution analysis was conducted using the MusiC R package v1.0^95^. First, the gene counts table from the bulk RNA-seq experiment was normalized and converted to counts per million (CPM). The gene expression profile matrix for each cell population, as defined by our single-cell analysis, was then used as input to estimate the proportion of each cell population within each bulk RNA-seq sample. To quantify the relative contribution of each differentially expressed genes upon *HSCHARME* KO to individual cell transcriptomes, and to cross-validate deconvolution analysis results, “KL-GSEA” was performed. We calculated per-cell Kullback-Leibler divergence scores comparing transcriptome expression profiles and a target distribution restricted to either genes downregulated (putative cardiac developmental genes) or upregulated in *HSCHARME* KO (bulk RNA-seq). KL scores for each cell were visualized using UMAP.

### Alternative Splicing Analysis

Cutadapt v3.2^96^ with parameters: *-u 1 -U 1 –trim-n* and Trimmomatic v0.39^97^ with parameters: *-PE ILLUMINACLIP:adapter_path:2:30:10:8:true LEADING:3 TRAILING:3 SLIDINGWINDOW:4:20* were used to remove adapter sequences and poor quality bases; for both software the minimum read length after trimming was set to 35. Then, STAR software v2.7.7a^81^ was used to align reads to GRCh38 genome using the following parameters: *-- outSAMstrandField intronMotif --outSAMattrIHstart 0 --outFilterType BySJout -- outFilterMultimapNmax 20 --alignSJoverhangMin 8 --alignSJDBoverhangMin 1 -- outFilterMismatchNmax 999 --outFilterMismatchNoverLmax 0.04 --alignIntronMin 20 -- alignIntronMax 1000000 --alignMatesGapMax 1000000 --outFilterIntronMotifs RemoveNoncanonical --peOverlapNbasesMin 3.* Alternative splicing events were identified using rMATS turbo version 4.1.1^48^, with the following parameters: *–variable-read-length, – allow-clipping, –readLength 150*, and *–libType fr-firststrand*. This analysis utilized the Ensembl gene annotation file (release 86) as the reference for splicing events. For downstream analyses, JCEC output from rMATS was selected. Splicing events were filtered based on false discovery rate (FDR) and Inclusion Level criteria. Only events with an FDR value of <0.05 and an Inclusion Level difference >0.1 or < -0.1 were considered significant and included for further examination (**Supplementary Table 3**). The rmats2sashimiplot Python script (available at https://github.com/Xinglab/rmats2sashimiplot) was used to visualize the selected alternative splicing events. This tool provided a graphical representation of splicing patterns, aiding in the interpretation of differential splicing events. Alternative splicing analysis in heart tissue comparing WT and *Charme* KO mice was performed following the same approach, using RNA-seq data available from the GEO dataset GSE200878^22^. To assess alternative splicing events conservation, the coordinates of exons regulated by *Charme* (either included or skipped) in mouse heart tissue were extracted and converted from mm10 genome to the hg38 human genome using the LiftOver tool^98^. The fraction of successfully lifted-over mouse-regulated exons that overlapped perfectly with regulated exons in human CM was then calculated, ensuring the consistency of the observed regulation between mouse and human datasets. To estimate the expected overlap by chance, 1,000 random samples of human exons were picked up from the ones detected by rMATS software (including not *pCHARME*-regulated). Then the overlap between human random exons and lifted-over mouse-regulated exons, generating a distribution of expected values.

### *pCHARME* sequence analysis

For the analysis of the k-mer composition, *pCHARME* was analyzed together with 1,987 genes annotated as “lncRNA” and expressed in WT CM (D20), each with an average FPKMs greater than 1. Gene annotations and transcript sequences were retrieved from Ensembl release 99^99^ using the biomaRt R package^100^. When multiple isoforms were associated with a gene, the longest isoform was selected to reduce sequence redundancy. The frequency of all possible combinations of seven nucleotides (7-mers) was computed for each long non-coding RNA. The *pCHARME* 7-mer signature was defined as the set of 7-mers exhibiting the highest frequency in *pCHARME* compared to other lncRNAs (top 0.1%). The sequence logo of the *pCHARME* 7-mers was generated using the Logomaker Python package v0.8^101^.

The RNA secondary structure of *pCHARME* was predicted using the RNAfold algorithm^59^ with standard parameters. Repeated and low complexity sequences were identified by scanning the transcript sequence with RepeatMasker v4.1.1^102^, using default settings and specifying the appropriate organism (Homo sapiens), with rmblastn v2.11.0+ as the internal search engine. The RepeatMasker output was subsequently converted to BED format using the RM2Bed.py utility provided by the software. To integrate the various analyses and visualize the results along the *pCHARME* sequence, the lncRNA was segmented from 5’ to 3’ into windows of 50 nucleotides in length, with sliding step of 10 nucleotides. The resulting bins were compiled into a BED file. BEDtools intersect v2.29.1^103^ was used to assign repeated elements to overlapping bins, while getfasta was applied to retrieve bin sequences, from which GC content and the occurrence of the *pCHARME* 7-mer signature were calculated. The Accessibility score was computed for each bin using the dot-bracket notation of the previously predicted *pCHARME* MFE structure, by calculating the fraction of unpaired nucleotides within each window. Finally, the ComplexHeatmap R package was used to visualize all results in a unified representation^104^.

### Analysis of Protein-RNA interactions

To identify RNA-binding proteins with high affinity for the *HSCHARME* transcript, we applied four complementary approaches: i. eCLIP-based motif enrichment analysis. We leveraged binding affinity scores of 150 RBPs for all possible 5-mers derived from the analysis of crosslinking induced truncations (CITS)^105^ in 223 eCLIP experiments^57^. For each experiment, the RBPs binding motifs were defined as the top 5 5-mers with highest PEKA scores from eCLIP datasets processed with Clippy or narrowPeaks softwares, or mCROSS^105^. To assess RBP affinity for *pCHARME*, we decomposed the *pCHARME*-specific 7-mers into 5-mers and selected RBPs for which the selected 5-mers matched the RBPs binding motifs in at least two analysis methods; ii. *in vitro* RNAcompete-derived binding profiles. We retrieved RBPs affinity scores (Z-scores) for all possible 7-mers from the RNAcompete dataset available in the CisBP-RNA database^60^. RBPs with *pCHARME* 7-mers ranked among their top binders (Z-score > 2) were selected as high-affinity candidates; iii. *in silico* sequence-based binding prediction with catRAPID. We used catRAPID software^61^ to predict interactions between full-length *pCHARME* and a reference library of RBPs. Proteins with a predicted interaction Z-score > 2 were considered high-affinity binders; iv. motif enrichment using catRAPID signature. We further identified candidate RBPs by evaluating the enrichment of known binding motifs within the *pCHARME* sequence using the motif analysis module of catRAPID. RBP binding enrichment analysis to pre-mRNA regions proximal to splicing events regulated by *pCHARME* was performed by focusing on exon inclusion and exclusion events upon its KO, as identified using the rMATS software. To map RNA-binding protein (RBP) binding sites, we used IDR peaks from the eCLIP datasets previously described^57^. To ensure compatibility with the eCLIP data, we first filtered the rMATS output from the *HSCHARME* KO versus WT comparison in CM, retaining only those genes also expressed in HepG2 and K562 cell lines (average FPKM > 1), which are the cell lines where eCLIP were performed. Next, for each exon significantly affected by *HSCHARME* loss (differentially spliced), we selected a matched control exon not affected by the lncRNA KO. Control exons were chosen with splicing junctions closely match the regulated exons in terms of splice junction counts (SJC) and intron junction counts (IJC), ensuring comparable expression levels. For each differentially spliced and control exon, we considered the following regions: the 3′ splice site (SS) of the exon upstream to the regulated one (defined as “upstream”); both the 5′ and 3′ SS of the regulated exon (defined as “center”), and the 5′ SS of the exon downstream to the regulated one (defined as “downstream”). For each site, 250 nucleotides of intronic sequence adjacent to the splice junction and 50 nucleotides of exonic sequence were analyzed, generating a total of 8 pre-mRNA region types: exonic-upstream, intronic-upstream, intronic-center(5’SS), exonic-center(5’SS), exonic-center(3’SS), intronic-center(3’SS), intronic-downstream and exonic-downstream.

Subsequently, BEDtools intersect was employed with the -s parameter to identify strand-specific overlaps between RBP binding sites, derived from each eCLIP experiment, and the pre-mRNA regions flanking dysregulated exons and their matched controls. For each eCLIP dataset, the number of regions either bound or unbound by the RBP was quantified separately for dysregulated and control exons, allowing the construction of a 2×2 contingency table to evaluate differential binding patterns. Then, Fisher’s exact test was applied to assess whether RBP binding was significantly enriched or depleted in the regions associated with dysregulated exons compared to controls. Finally, Benjamini-Hochberg false discovery rate (FDR) correction was used to adjust p-values for multiple hypothesis testing.

### Pathological dataset analysis

Pathological datasets raw data (FASTQ files) were retrieved from Gene Expression Omnibus (GEO) database with the following accession IDs: GSE130036 for HCM and GSE116250 for DCM. Data were pre-processed and analyzed as described in the RNA-Sequencing analysis section (**Supplementary Table 5**). Intersections between *HSCHARME* KO and the pathological datasets were performed using a 0.05 adjusted p-value cutoff for all genes in all the analyzed datasets and reporting the log2 fold change values for each gene. Figures were prepared with Prism 7.0 software and edited with Adobe Illustrator 2024.

### Flow cytometry assay

For PDGFRA, CD56 and CD82 cell surface markers, CM were washed twice with DMEM F12 and dissociated as single cells using 0.25% trypsin-EDTA. Cells were stained with the diluted (1:10) primary antibody (CD82, see **Supplementary Table 6**) for 45 minutes in Flow Cytometry Buffer (FCB; 5% FBS in PBS). After two washes in FCB, cells were incubated for 30 min at RT with a secondary antibody (Alexa Fluor 488, Life Technologies) at 1:300 dilution or with a conjugated antibody (APC-PDGFRA 1:20 and 488-CD56 1:20) in FCB. After two washes, stained cells were analyzed by CytoFLEX SRT Cell Sorter (Beckman Coulter). Data were collected from at least 10.000 events and analyzed using Kaluza software (Beckman Coulter). For TNNT2, were fixed in 4% paraformaldehyde (PFA) in PBS for 15 minutes at room temperature (RT) before staining. After fixation, cells were washed twice by being resuspended in Flow Cytometry Buffer (FCB; 5% FBS in PBS), centrifuged for 5 min at 250 g and the supernatant was discarded. A third wash with FCB-0.2% Triton X-100 was performed as described above. Cells were stained with the diluted (1:100) primary antibody (TNNT2, see **Supplementary Table 5**) in FCB-0.2% Triton X-100 for 45 min at RT. After two washes in FCB, cells were incubated for 30 min at RT with a secondary antibody (Alexa Fluor 488, Life Technologies) at 1:400 dilution, washed twice more and resuspended in PBS. Stained cells were analyzed and sorted on a flow cytometer (BD Bioscience, Milan, Italy). Data was collected from at least 10,000 events. Flow cytometry data were analyzed with Flowing software 2.5.1. After exclusion of debris and doublets based on light scatter properties, single cells were analyzed for the expression of TNNT2 and reported as the percentage of marker-positive cells or as mean fluorescence intensity.

### Immunofluorescence and RNA-FISH assays

Glass coverslips were prepared as for ^106^ by treatment with poly-L-lysine (50 mg/ml in water) for 30 minutes under UV light and overnight treatment with Laminin (2 mg/ml in water) at 37°C. After 20 days of differentiation, hiPSC-derived CM were washed twice with DMEM F12 and dissociated as single cells using 0.25% trypsin-EDTA. Isolated CM were re-plated on the treated coverslips in RPMI 1640 Medium (Life Technologies) supplemented with B27 Supplement and Rock inhibitor (1:1000). After 24 hours, cells were fixed in 4% paraformaldehyde (PFA) in PBS for 20 minutes at 4°C prior staining. For immunofluorescence, cells were permeabilized with 0.2% Triton X-100 in PBS for 20 minutes at room temperature (RT) and blocked with 1% Goat Serum (GS) in PBS for 20 minutes at RT. The primary antibody (TNNT2, see **Supplementary Table 6**) was diluted (1:200) in 1% GS-PBS and incubated overnight at 4°C. The cells were rinsed with PBS and incubated for 1 hour with a secondary antibody (Alexa Fluor 488, Life Technologies) at 1:300 dilution, rinsed, counterstained with DAPI for 5 minutes at RT and mounted on cover-glasses.

For RNA-FISH experiment, staining was carried out with the HCR RNA-FISH technology according to manufacturer protocol (https://www.molecularinstruments.com/hcr-rnafish), with a specific set-up thought for lncRNA staining. Images were acquired on Carl Zeiss Axio Vert.A1 Microscope or with an inverted confocal Olympus IX73 microscope equipped with a Crestoptics X-LIGHT V3 spinning disk system and a Prime BSI Express Scientific CMOS camera. The acquisitions were obtained using a 20X air objective (IF-only samples) or UPLANSApo 60X (NA 1.35) oil objective (FISH/IF samples) and collected with the MetaMorph software (Molecular Devices). SC35 domains/*pCHARME* and PTBP1/*pCHARME* overlapping quantification was performed by ImageJ tolls in JACoP package in order to obtain a Mander’s overlap coefficient (MOC) from single ROIs of original confocal images.

### Cross-linking immunoprecipitation (CLIP) assay

hiPSC-CM (D20) were washed twice with PBS and UV-crosslinked (4000 μJ) in a Spectrolinker UV Crosslinker (Spectronics Corporation). Cells were harvested in lysis buffer (20 mM Tris-HCl, 100 mM NaCl, 0.5 mM EDTA, 0.5% [v/v] NP40, 0.1% SDS, 0.5 mM DTT, 1× PIC), incubated on ice for 15 min and sonicated at low-intensity four times with Bioruptor Plus sonication device to ensure membrane lysis. Lysate was diluted to a final concentration of 1 mg/ml and PTBP1 CLIP was carried out as in ^107^

### Morphological analysis

For morphological analysis, 10 cells for each field were individually selected and analysed using ImageJ standard tools. 5 fields were considered for each biological replicate. Cell area and circularity were computed with the ImageJ measure tool with circularity=1 indicating a perfect circle. Elongation represents the ratio of the major axis length to the minor axis length, as described in ^42^. For each parameter, the average of each biological replicate was represented as a dot in the plots. Statistical analyses were performed on the overall average measure of the 5 biological replicates.

### Cardiomyocytes contraction analysis

Spontaneous CM contraction was assessed after 20 days of differentiation. To quantify beat rate and contraction dynamics, light brightfield videos of WT and ΔP beating monolayers were recorded with an OkoLab stage top incubator equipped with 20X air objectives under controlled CO₂, pH and temperature conditions. One video per well was taken across n=5 biological replicates. Video recordings were processed using a MuscleMotion macro^108^ in FIJI/ImageJ software (v2.9.0). MUSCLEMOTION outputs included contraction duration, time to peak and peak-to-peak time.

### Statistical methods and rigour

For RT-qPCR and flow cytometry experiments, statistical analyses were performed on log₂-transformed fold change (LogFC) values to stabilise variance and approximate a normal distribution using a two-tailed one-sample t-test against the null hypothesis of zero (no change). For experiments with n=5, data were assessed for normal distribution with a Shapiro-wilk test, and a parametric/non-parametric statistical test was used accordingly (Student’s t-test when data are normally distributed and Wilcoxon-Mann-Whitney when they are not). Details regarding statistical tests, p-values, and sample sizes (n) are provided in the corresponding figure legends.

## Data availability

The data presented in this study will be openly available in NCBI Gene Expression Omnibus (GEO) database (https://www.ncbi.nlm.nih.gov/geo/) with reference number GSE273100.

## Acknowledgments

The authors acknowledge Prof. Alessandro Rosa for kindly providing the WT1 hiPSCs, Dr. Davide Mariani and the Genomic Facility of the Center for Human Technologies of IIT for support in RNA-sequencing experiments, Dr. Marcella Marchioni for technical help with cell culture and Dr. Maria Celardo and Marco Simula for helpful discussion. This research was funded by: 1) Sapienza University (RM12117A5DE7A45B and RM123188F6B80CE4), to M.B.; 2) Consiglio Nazionale delle Ricerche-CNR (projects DBA.AD005.225-NUTRAGE-FOE2021 and DSB.AD006.371-InvAt-FOE2022), to P.L.; 3) Avvio alla Ricerca Type 2 (Prot. AR223188B40CB2D0) to G.B.; 4) European Union - NextGenerationEU: National Center for Gene Therapy and Drug based on RNA Technology, CN3 - code: CN00000041; National Recovery and Resilience Plan (NRRP) MUR - M4C2 - Action 1.4 - Call “Potenziamento strutture di ricerca e di campioni nazionali di R&S” (Spoke 3 “Neurodegeneration”, CUP: B83C22002870006, to M.B. and Spoke 6 “RNA Drug Development”, CUP B83C22002860006, to P.L.); 5) by the European Union - Next-GenerationEU - National Recovery and Resilience Plan (NRRP) - M4C2, INVESTMENT N. 1.1, Call “PRIN 2022” (project 2022BYB33L, CUP: B53D23016090006), to M.B. and P.L.; 6) by the European Union - Next-GenerationEU - National Recovery and Resilience Plan (NRRP) - M4C2, INVESTMENT N. 1.1, Call “PRIN 2022 PNRR” (project P2022FFEWN RNA2FUN, CUP: B53D23026140001), to M.B.

## Author contributions

M.B. designed, conceived and acquired funding for the study; M.B. and P.L. secured fundings; G.B developed the PA and RHE CRISPR-Cas9 edited cell lines and performed all the gene expression, morphological and functional analyses in hiPSC-derived cardiomyocytes; F.D. developed the *Δ*P cell line and provided support for the gene expression analyses; A.S. performed the alternative splicing analysis on the WT, *Δ*P and GSE200878 dataset, as well as the conservation and *in silico* interaction analyses; A.P. performed the computational analyses on the WT, *Δ*P, GSE130036 and GSE116250 datasets; A.D. carried out computational analyses on the syn7818379 datasets and the cellular deconvolution analysis; G.S. provided support with the hiPSC handling and gene expression analyses; T.S. performed the RNA-FISH experiments; D.T. performed the flow cytometry assay. The project was supervised and written by M.B. with major contributions from G.B. and P.L. and input from the other authors.

## SUPPLEMENTARY FIGURES

**Supplementary Fig. 1.**
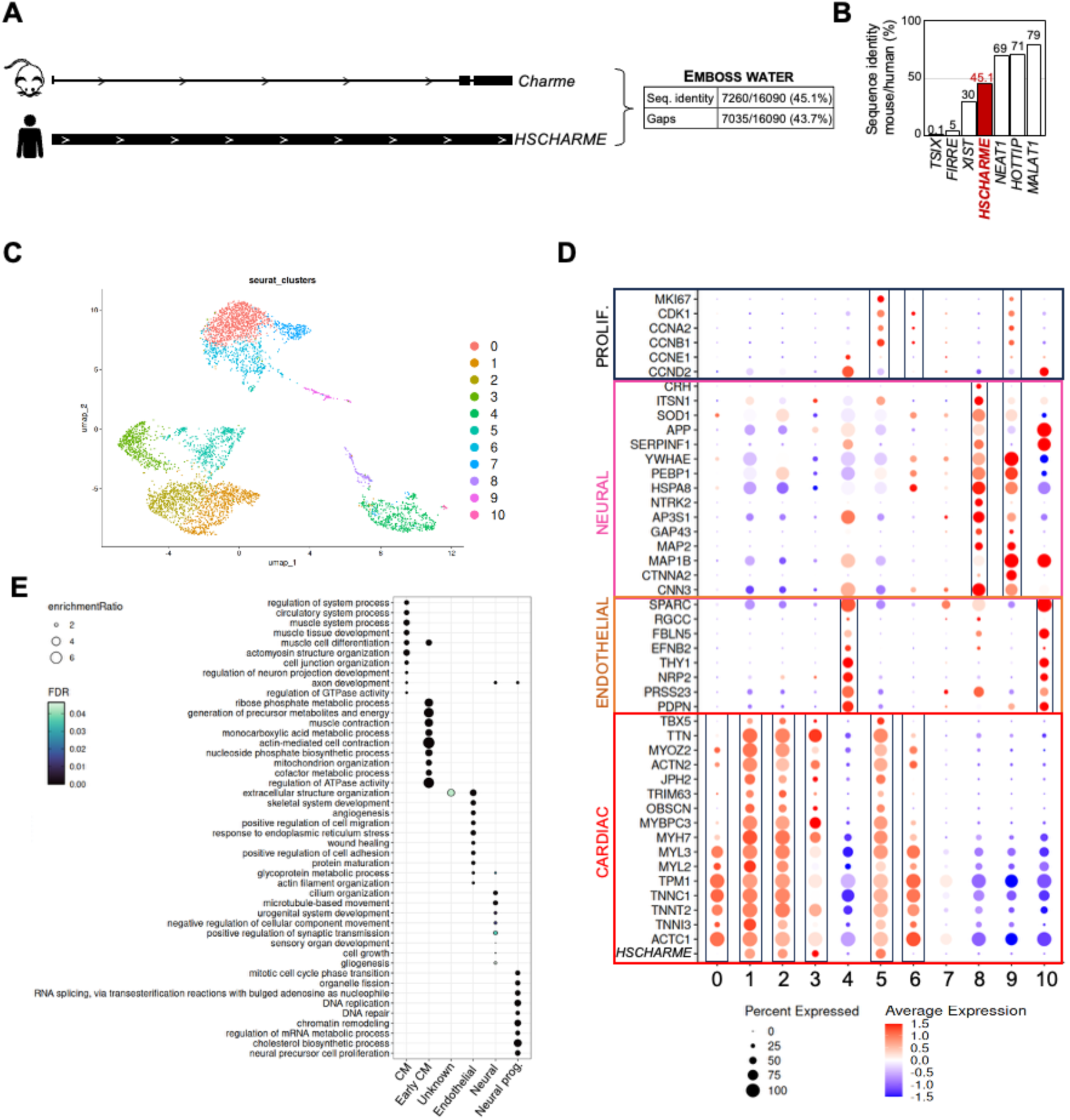
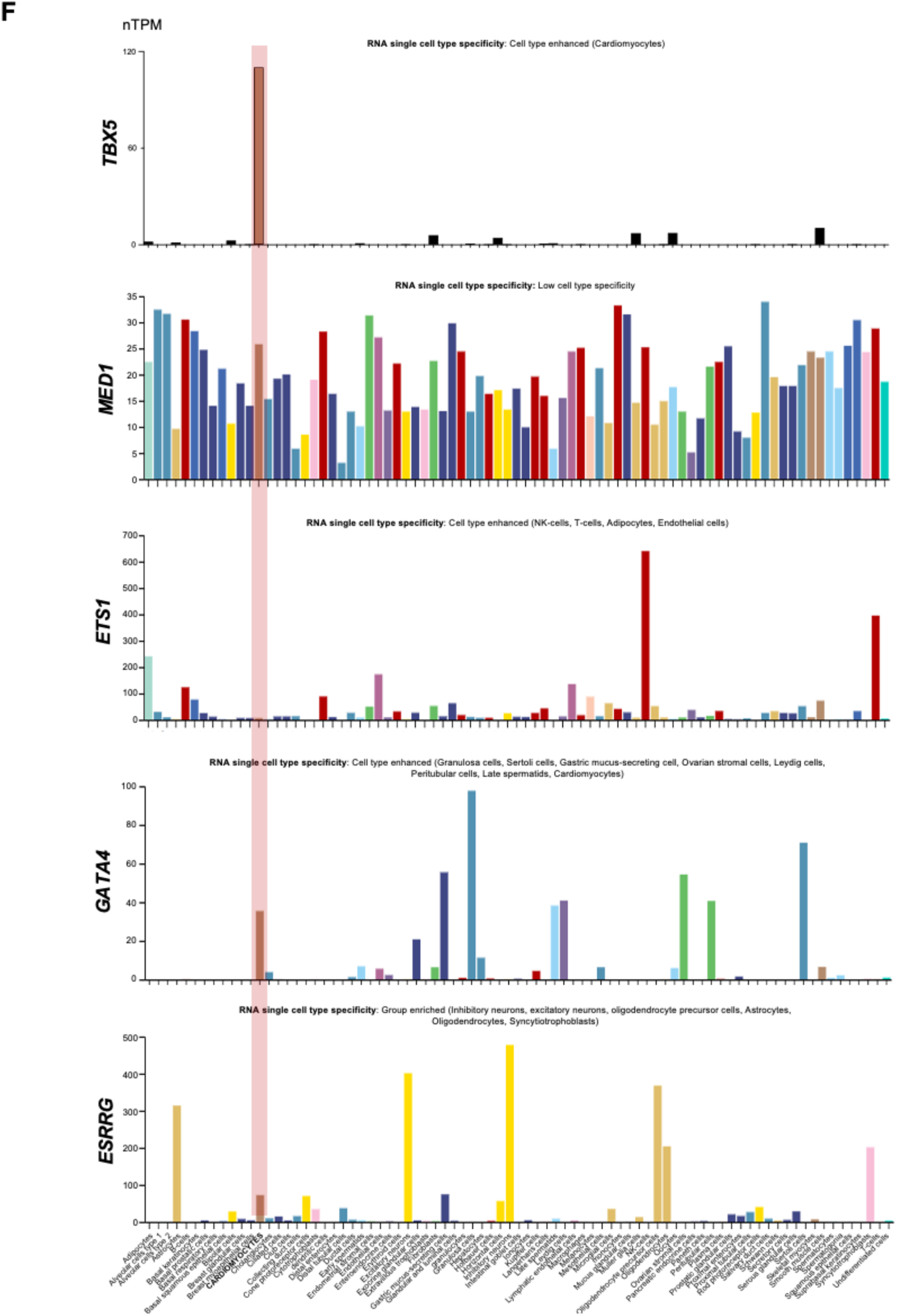
**A)** Table reports the sequence conservation values between the mouse and human *Charme* genes. The data were obtained by local sequence using the Smith– Waterman algorithm available at Emboss water (https://www.ebi.ac.uk/jdispatcher/psa/emboss_water). **B)** Barplot reporting the percentage of sequence identity between mice and humans for a subset of well-known lncRNAs. C) UMAP visualization of single-cell transcriptomic profiles from hiPSC-CM (2857 cells from Day 14, 2321 cells from Day 45 as in ^31^. Cells are colored based on the clusters identified through unsupervised clustering by Seurat. **D)** Dot plot of scaled average gene expression across cell clusters of neural, endothelial, cardiac and proliferative cell markers. Dot size corresponds to the proportion of cells within a cluster expressing the indicated gene, and color indicating the scaled expression level. Boxes indicates the correspondence between gene markers and cell types. **E)** Dot plot of gene ontology categories across cell populations. The plot displays only the weighted set categories. Dot size corresponds to the enrichment ratio of terms, and color indicates the statistical significance (FDR). **F)** Quantification of *ESRRG, MED1, ETS1, GATA4 and TBX5* expression levels at single cell resolution as reported in the Human Protein Atlas Portal. RNA expression is quantified in nTPM (nTPM = normalized Transcripts Per Million).

**Supplementary Fig. 2.**
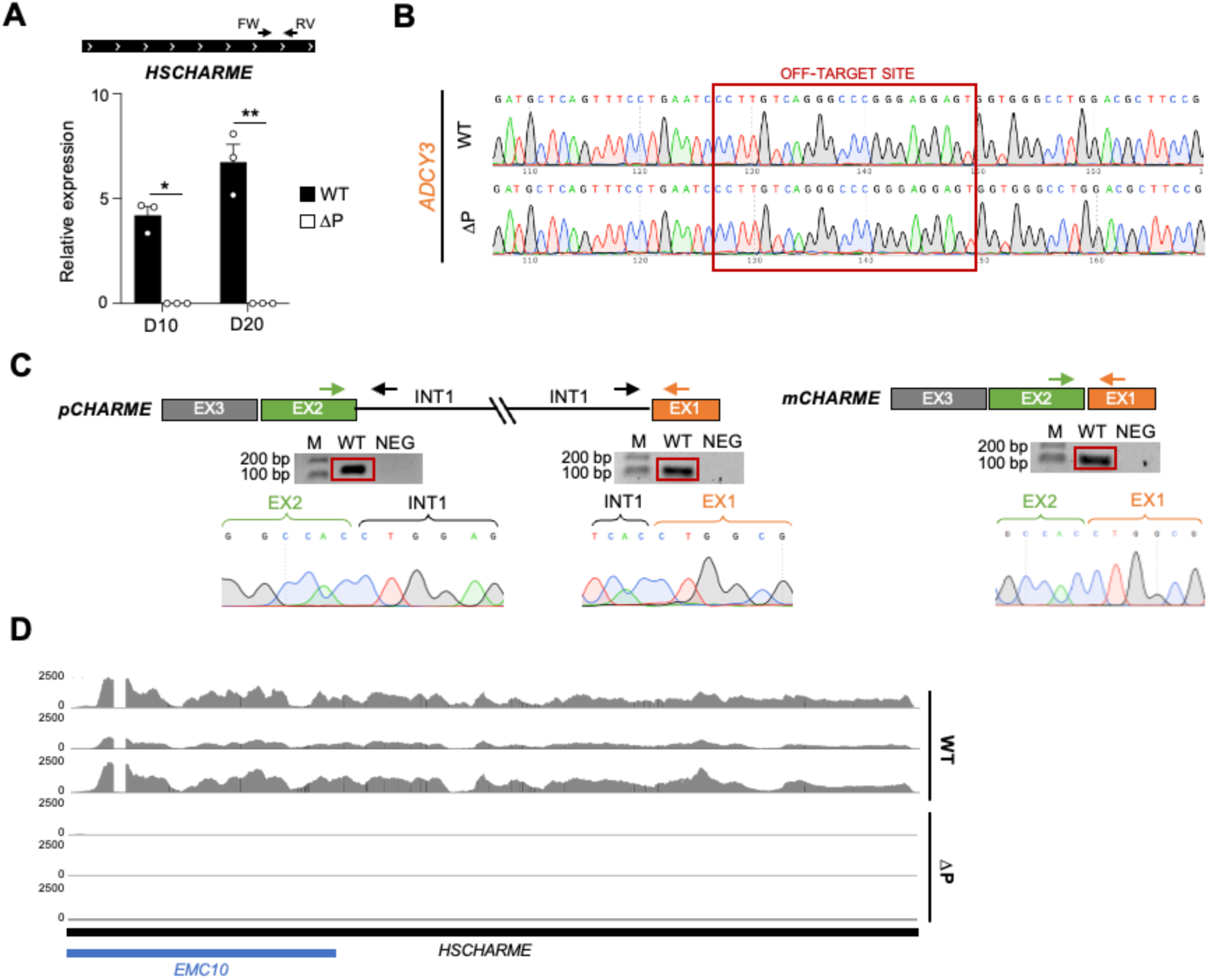
**A)** RT-qPCR amplification of *HSCHARME* from WT vs ΔP D10 and D20 CM. Data were normalized to *ATP5O* mRNA and represent relative expression means (2^^-DCt^) ± SEM of 3 biological replicates. Statistical tests were conducted on log₂-transformed fold change (LogFC) values compared to the control sample (WT). Data information: *p < 0.05; ***p < 0.01; ****p<0.0001; one-sample two-tailed Student’s t-test against the null hypothesis of zero (no change). **B)** Representative example of sequencing results from WT and ΔP on the predicted off-target *ADCY3* gene. The red square indicates the region interested in the off-targeting. **C)** sqRT-PCR amplifications of *pCHARME* (5’ and 3’-end) and *mCHARME* from WT-CM. Bands in the red square were extracted and subjected to Sanger sequencing as reported below. **D)** RNA-seq normalized read coverage tracks (TPM) across *HSCHARME* locus from WT and ΔP-CM (D20).

**Supplementary Fig. 3.**
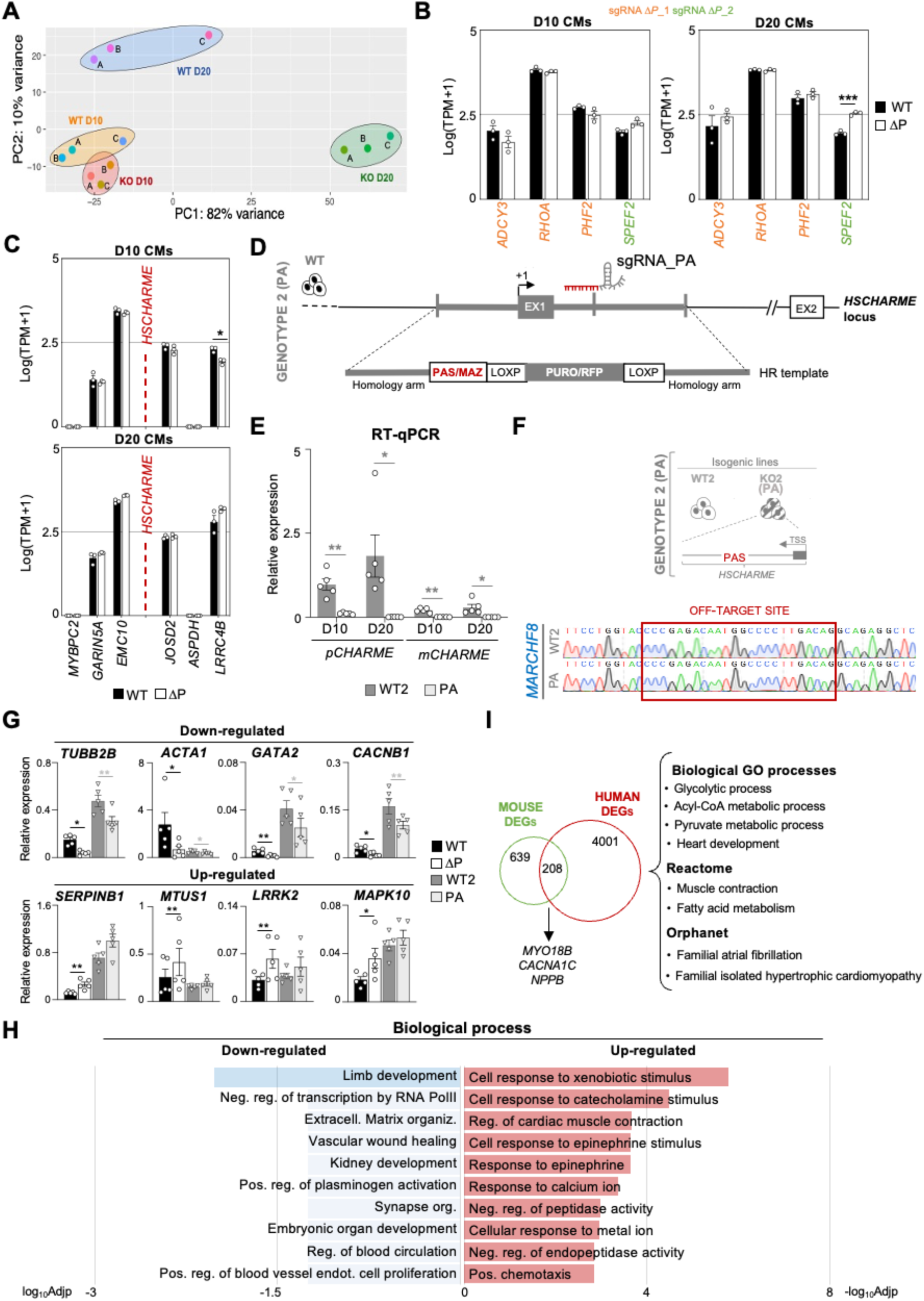
**A)** Principal Component Analysis (PCA) performed on WT and ΔP RNA-seq datasets at D10 and D20 hiPSC-derived CM. **B)** Quantification by RNA-seq (TPM) of sgRNAs off-targets expression in WT vs ΔP hiPSC-derived D10 (left panel) and D20 (right panel) CM. Data are represented as Log(TPM+1) to avoid negative values. Data information: ***p<0.001, FDR. **C)** Quantification by RNA-seq (TPM) of *HSCHARME*-neighbouring genes expression in WT vs ΔP hiPSC-derived D10 (upper panel) and D20 (lower panel) CM. Data are represented as Log(TPM+1) to avoid negative values. **D)** Representation of the genome editing strategy design followed to obtain the PA hiPSC cell line by CRISPR/Cas9. **E)** RT-qPCR amplification of *pCHARME* and *mCHARME* from WT2 *vs* PA (D10 and D20) CM. Data were normalized to *ATP5O* mRNA and represent relative expression means (2^^-DCt^) ± SEM of 5 biological replicates. Statistical tests were conducted on log₂-transformed fold change (LogFC) values compared to the control sample (WT2). **F)** Representative example of sequencing results from WT2 and PA on the predicted off-target *MARCHF8.* The red square indicates the region interested in the off-targeting. A schematic representation of the genome editing strategy is represented on the top. **G)** RT-qPCR quantification of down-regulated (upper panel) and up-regulated (lower panel) DEGs in WT *vs* ΔP (black) and WT2 vs PA (grey) hiPSC-derived CM (D10). Data were normalized to *ATP5O* mRNA and represent relative expression means (2^^-DCt^) ± SEM of 5 biological replicates. Statistical tests were conducted on log₂-transformed fold change (LogFC) values compared to the control sample (WT for ΔP-CM and WT2 for PA-CM). **H)** Gene ontology (GO) enrichment analysis performed with EnrichR on down-regulated (left) and up-regulated (right) DEGs in WT *vs* ΔP hiPSC-derived CM (D10). Bars indicate the +/–log_10_ adjusted p-value (log_10_Adjp and –log_10_Adjp) of the top enriched biological processes. Categories represented by darker bars show an Adjp <0.05. **I)** Venn diagrams depicting the intersection between mouse and human DEGs in response to *Charme* ablation. The top GO categories for biological process, Reactome and Orphanet are shown on the right. Notable genes are reported below. Data information: *p < 0.05; ***p < 0.01; ****p<0.0001; one-sample two-tailed Student’s t-test against the null hypothesis of zero (no change).

**Supplementary Fig. 4.**
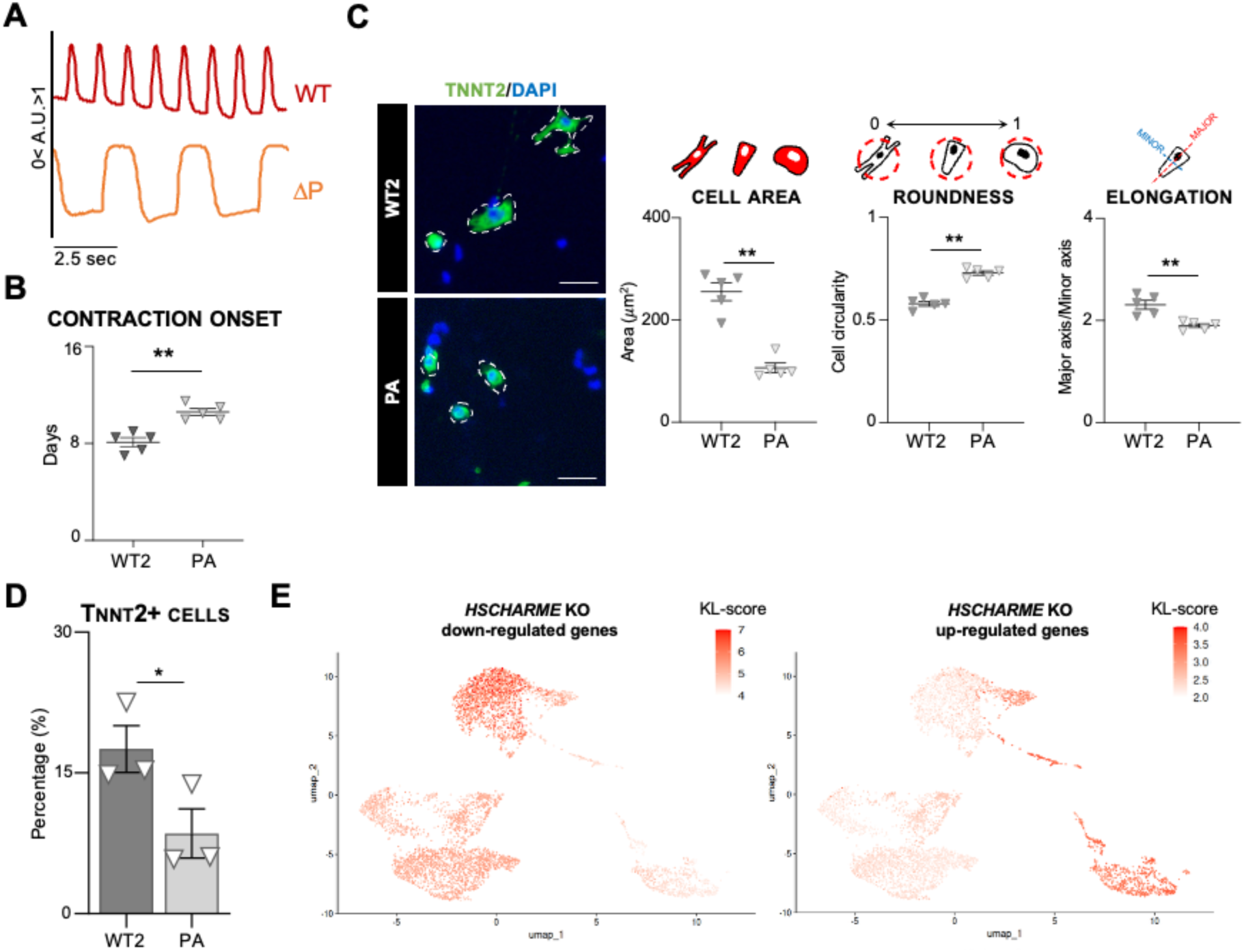
**A)** Representative traces of normalized spontaneous contraction profile of WT (red) and ΔP (orange) hiPSC-derived CM (D20). Traces derive from the analisys of brightfield 2D videos with the MUSCLEMOTION plugin. **B)** Onset (days) of spontaneous contraction of WT and ΔP hiPSC-derived CM. Data represent means ± SEM of 5 biological replicates. **C)** Left panel: Representative images of WT2 and PA hiPSC-derived CM (D20) selected for morphological analyses (white dashed outlines) after TNNT2 (green) and DAPI (blue) staining. Scale bar=50 µm. Right panel: Population measurements of CM morphological features. A schematic representation of the specific measure quantified is shown above each plot. Black bars represent means ± SEM of 5 biological replicates. **D)** Flow cytometry quantification of cardiac troponin-T (TNNT2) positive cells (%) in WT2 and PA hiPSC-derived CM (D20). Data represent mean ± SEM of 3 biological replicates. Data information: *p < 0.05; one-sample two-tailed Student’s t-test on LogFC against the null hypothesis of zero (no change). **E)** UMAP visualization of KL (Kullback–Leibler) score distribution across single cells (see **“Material and Methods”** for details). The left panel (“KL Score Down”) displays the KL enrichment score of downregulated genes in *HSCHARME* ΔP at day 20. The right panel (“KL Score Up”) displays the KL enrichment score of upregulated genes in *HSCHARME* ΔP (D20). Data information: *p < 0.05, **p < 0.01, ***p < 0.001, paired two-tailed Student’s t-test.

**Supplementary Fig. 5.**
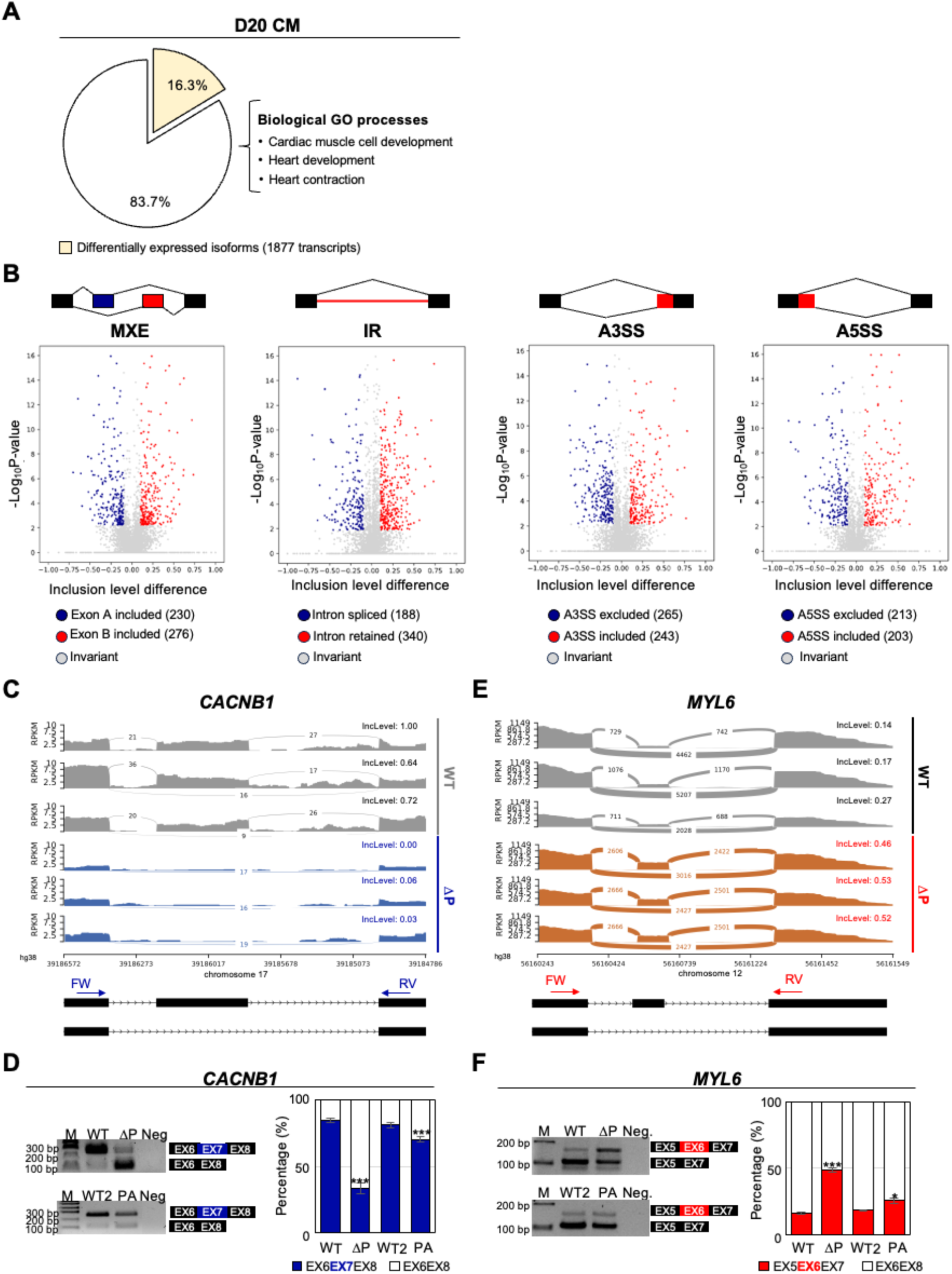
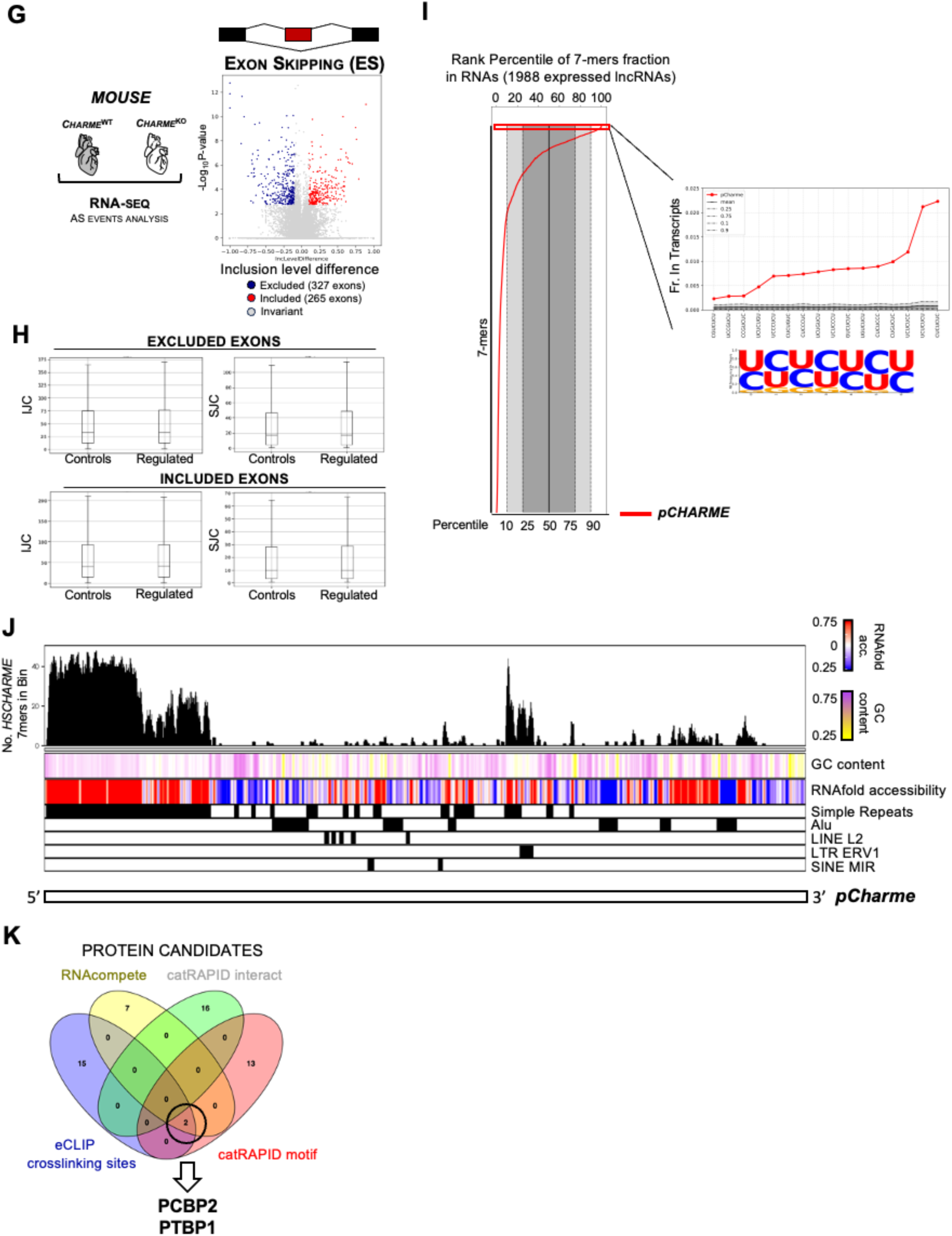
**A)** Pie Chart representing the fraction of significant (FDR<0.05) differentially expressed transcript isoforms in ΔP *vs* WT CM (D20). The top 3 GO categories for biological process and molecular function are shown on the right. **B)** Volcano plots depicting significantly altered splicing events (FDR<0.05; Inclusion level variations stronger than 10%) in ΔP-CM (D20). All classical splicing alterations are reported, such as Exon skipping (ES); Intron retention (IR); alternative 3’ splice site (A3SS); alternative 5’ splice site (A5SS) and Mutually exclusive exons (MXE). X-axis represent inclusion ratio while y-axis represent –log_10_ of P-value. A schematic representation of the specific investigated event is shown above each plot. **C)** Sashimi plots of *CACNB1* exon skipping events in WT and ΔP hiPSC-derived CM (D20). The X-axis indicates genomic location while the Y-axis reports expression levels as Reads Per Kilobase of transcript per Million mapped reads (RPKM). The lines connecting different exons (bridges) indicate reads on the specific splicing junction. Quantification of splicing reads is reported in each bridge while Inclusion levels (IncLevel) are reported above each sample. The exonic structure of the event is shown below together with oligonucleotides position used for sqRT-PCR validation. **D)** Left panel: Representative electrophoresis analyses of *CACNB1* exon skipping events by sqRT-PCR from WT *vs* ΔP or WT2 *vs* PA CM (D20). Right panel: Quantification of sqRT-PCR amplifications calculated by the intensity of the exon-including and the exon-excluding band and plotted as percentage. Each plot reports the mean percentage ± SEM of 5 biological replicates. **E)** Sashimi plots of exon skipping events in the *MYL6* transcripts in WT and ΔP hiPSC-derived CM (D20). The X-axis indicates genomic location while the Y-axis reports expression levels as Reads Per Kilobase of transcript per Million mapped reads (RPKM). The lines connecting different exons (bridges) indicate reads on the specific splicing junction. Quantification of splicing reads is reported in each bridge while Inclusion levels (IncLevel) are reported above each sample. The exonic structure of the event is shown below together with oligonucleotides position used for sqRT-PCR validation. **F)** Left panel: Representative electrophoresis analyses of sqRT-PCR for the exon skipping event in *MYL6* in WT vs ΔP and WT2 vs PA D20 CM. Right panel: Quantification of sqRT-PCR amplifications calculated by the intensity of the exon-including and the exon-excluding band and plotted as percentage. Each plot reports the mean percentage ± SEM of 5 biological replicates. **G)** Volcano plot depicts significant (FDR<0.05; Inclusion level variations stronger than 10%) ES events in murine *Charme* WT and KO post-natal (PN) hearts. X-axis represents exon inclusion ratio, y-axis represents –log10 of P-value. A schematic representation of the investigated event is shown above. **H)** Boxplots display the distribution of Intron Junction Counts (IJC) and Splice Junction Counts (SJC) for exon exclusion events (upper panel) and exon inclusion events (lower panel) following *pCHARME* depletion, along with their corresponding matched control events. Statistical significance was assessed using Mann-Whitney U-test. All the reported comparisons showed no significant differences (p-value > 0.05). **I)** Left panel: Percentile rank distribution of 7-mer frequencies in *pCHARME* compared to a reference set of 1,987 lncRNAs expressed in CM. Each point on the red line represents the rank percentile of an individual 7-mer based on its frequency in *pCHARME* relative to the lncRNA reference set. The x-axis denotes the percentile scale, while the y-axis refers to each of the possibile 7-mer. The light-gray shaded area represents the 10th–90th percentile range of the background distribution, and the dark-gray area highlights the interquartile range (25th–75th percentiles). The median (50th percentile) is marked by a black line. Right panel: Absolute frequency of each of the 16 *pCHARME*-specific 7-mers (red line) compared to their distribution observed in the background lncRNAs. Gray areas again denote the 10th–90th (light) and 25th–75th (dark) percentile bands, with the mean value across background lncRNAs shown in black. The sequence logo generated from the 16 *pCHARME*-specific 7-mers, illustrating their consensus UC-rich composition, is shown below. **J)** Heatmap showing key features of *pCHARME* sequence. The transcript is represented from the 5’ end (left) to the 3’ end (right) in consecutive 50-nucleotide segments with a 10-nucleotide sliding step. The top panel displays a bar plot reporting the total number of *pCHARME*-specific 7-mer occurrences within each segment. The heatmap below illustrates three distinct features along the transcript. The first is the GC content where segments with higher GC content are shown in violet, while those with lower GC content appear in yellow. The second feature is RNA accessibility, computed using RNAfold software and reported as the fraction of unpaired nucleotides in each segment; segments with higher accessibility are colored in red, and those with lower accessibility in blue. The third feature is the annotation of repetitive elements, computed using RepeatMasker software, where each segment is marked in black to indicate the presence of the specified repeat elements, or in white if absent. **K)** Venn Diagram depicting the overlap among proteins predicted to interact with *pCHARME* sequence using RNAcompete, eCLIP crosslinking sites, catRAPID interact and catRAPID motif approaches (see **“Material and Methods”** for details).

**Supplementary Fig. 6.**
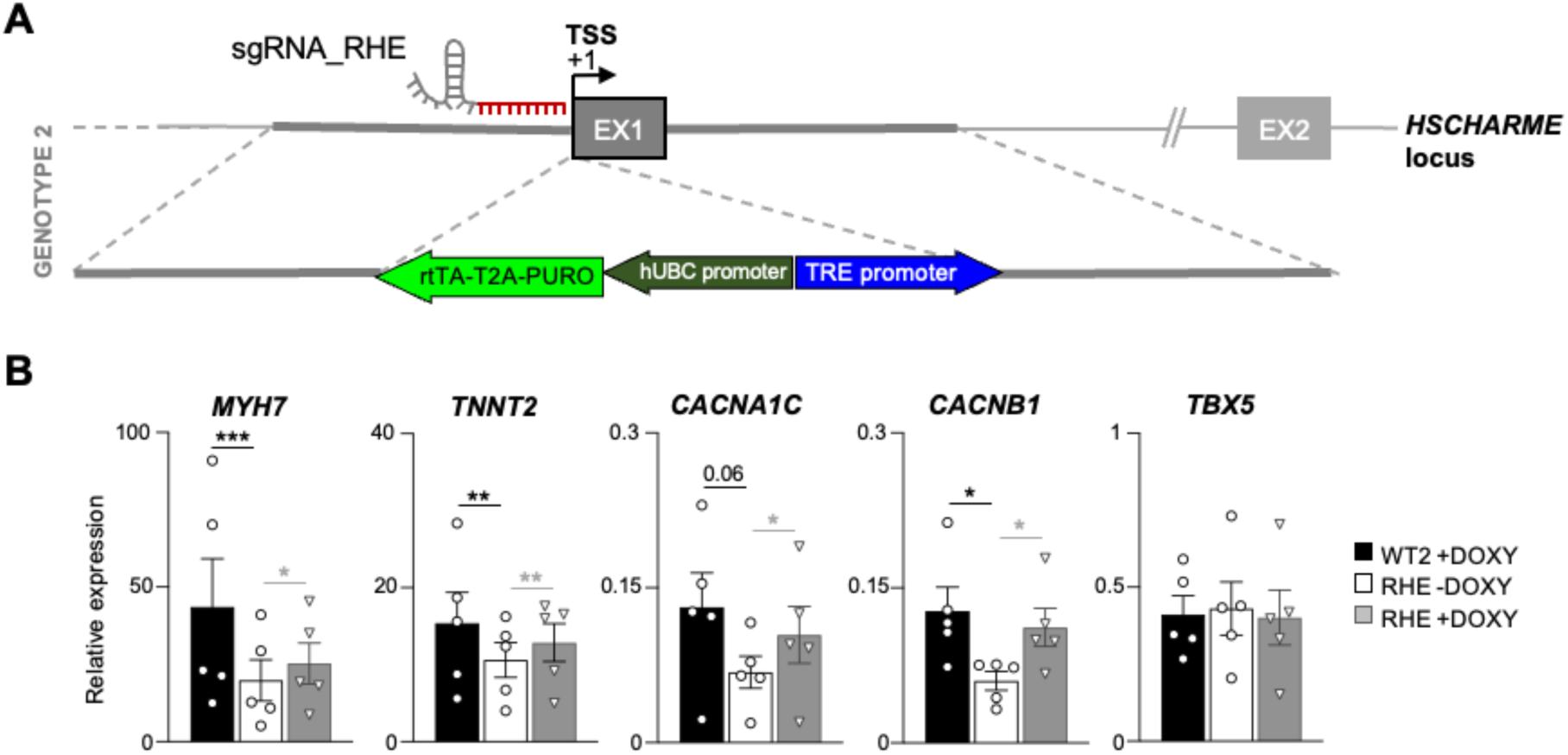
**A)** Schematic representation of the CRISPR/Cas9 genome editing design used to obtain the *RHE* hiPSCs. **B)** RT-qPCR quantification of selected mRNA in WT2, RHE (-DOXY) and RHE (+DOXY) hiPSC-derived CM (D20). Data were normalized to *ATP5O* mRNA and represent relative expression means (2^^-DCt^) ± SEM of 5 biological replicates. Statistical tests were conducted on log₂-transformed fold change (LogFC) values compared to the control sample (RHE -DOXY). Data information: *p < 0.05; ***p < 0.01; ****p<0.0001; one-sample two-tailed Student’s t-test against the null hypothesis of zero (no change).

